# Identification of *V. parvula* and *S. gordonii* adhesins mediating co-aggregation and its impact on physiology and mixed biofilm structure

**DOI:** 10.1101/2024.07.16.603716

**Authors:** Louis Dorison, Nathalie Béchon, Camille Martin-Gallausiaux, Susan Chamorro-Rodriguez, Yakov Vitrenko, Rania Ouazahrou, Romain Villa, Julien Deschamps, Romain Briandet, Simonetta Gribaldo, Jean-Marc Ghigo, Christophe Beloin

## Abstract

The dental plaque is a polymicrobial community where biofilm formation and co-aggregation, the ability to bind to other bacteria, play a major role in the construction of an organized consortium. One of its prominent members is the anaerobic diderm *Veillonella parvula,* considered as a bridging species, which growth depends on lactate produced by oral *Streptococci*. Understanding how *V. parvula* co-aggregates and the impact of aggregation has long been hampered due to the lack of appropriate genetic tools. Here we studied co-aggregation of the naturally competent strain *V. parvula* SKV38 with various oral bacteria and its effect on cell physiology. We show that *V. parvula* requires different trimeric autotransporters of the type V secretion system to adhere to oral *Streptococci* and *Actinomyces*. In addition, we describe a novel adhesin of *Streptococcus gordonii,* VisA (SGO_2004), as the protein responsible for co-aggregation with *V. parvula*. Finally, we show that co-aggregation does not impact cell-cell communication, which is mainly driven by environmental sensing, but plays an important role in the architecture and species distribution within the biofilm.

## INTRODUCTION

Bacterial attachment to other bacteria is a key step in the formation of bacterial biofilm. This adhesion is termed auto-aggregation when the adhesion occurs with a genetically identical bacteria and co-aggregation when different species or strains are involved. While auto-aggregation is known to enhance stress resistance, antibiotic tolerance, and virulence, the specific role of co-aggregation remains largely understudied^1^, except in the contexts of the dental plaque and certain aquatic environments^2–4^.

The dental plaque is an important polymicrobial biofilm whose perturbation can lead to the development of caries and periodontitis^5,6^. The formation of the dental plaque is a stepwise process which begins with the adhesion to the teeth surface of early colonizers comprised of oral streptococci, including *Streptococcus gordonii, S. oralis* and *S. mitis* and *Actinomyces spp.*. Then, bridging species such as *Veillonella* and *Fusobacterium* co-aggregate with the early colonizers forming an adhesion substrate for late biofilm commensal colonizers but also the opportunistic pathogens *Porphyromonas gingivalis*, *Treponema denticola* and *Tannerella forsythia*^2^. Co-aggregation is mostly driven by adhesins ^7–12^, few of which have been identified, including *P. gingivalis* major and minor fimbriae^13,14^, which interacts with *S. gordonii* SspB adhesin and GADPH, and the *F. nucleatum* autotransporters RadD and Fap2 ^9,11,15^. However, most of the molecular actors of oral biofilm co-aggregation mechanisms are currently unknown.

*Veillonella* are strict anaerobic diderm firmicutes and seven *Veillonella* species can be found in the dental plaque^16^ where they rely on lactate produced by oral streptococci as a carbon source^17^. Oral *Veillonella* species possess extensive aggregative properties contributing to their colonization of the oral environment ^7^ in which the physical proximity resulting from aggregation with their different partners likely facilitates their metabolic integration in the oral biofilm. For instance, *V. parvula* (previously *V. atypica*) strain PK1910 induces the expression of the *S. gordonii* amylase *amyB* in a distance-dependent manner, possibly to increase lactic acid production^18,19^. *V. atypica* was also shown to produce a catalase protecting *F. nucleatum* from reactive oxygen species produced by *S. gordonii*^20^.

While *Veillonella* adhesive properties have been first characterized more than 30 years ago ^21,22^, the underlying molecular actors of co-aggregation and its physiological consequences remained elusive until recently. Indeed, it was recently shown that *V. atypica* OK5 possesses eight trimeric autotransporter adhesins (TAA) belonging to the type Vc secretion system family. One of them, Hag1, mediates adhesion to oral bacteria and buccal cells^23^. On the other side, several oral *Veillonella* species, including *V. atypica* OK5, co-aggregate with *S. gordonii* Hsa adhesin^24^. However, a more extensive mechanistic characterization of the *Veillonella* adhesin repertoire was hampered due to the lack of genetic tools described for this genus. *V. parvula* strain SKV38 is a recently described naturally competent isolate that is readily genetically engineered^25^. We have recently shown that it possesses nine TAAs, named VtaA to -I, and 3 classical monomeric autotransporters, named VmaA to -C. Both VtaA and a gene cluster coding for 8 TAA adhesins were shown to be important for surface adhesion and biofilm formation^25^.

Here, we investigated the capacity of *V. parvula* SKV38 to co-aggregate with common oral bacteria and studied the physiological impact of this co-aggregation. We found that, in addition to mediating auto-aggregation, VtaA is also involved in co-aggregation with *S. oralis* while two other adhesins encoded in an adhesin cluster, VtaE and VtaD, are involved in co-aggregation with *S. gordonii* and *Actinomyces oris*. We also identified a novel adhesin of *S. gordonii*, VisA (SGO_2004), as the possible interacting partner of *V. parvula* VtaE/VtaD. Analysis of the transcriptomic profiles of both bacteria in coculture with or without aggregation suggested a very limited impact of aggregation on gene expression. Furthermore, we showed that absence of co-aggregation results in spatial segregation of the two species biofilms, suggesting that co-aggregation would be necessary to generate the architecture of a healthy dental plaque biofilm. In conclusion, this study contributes to provide a better mechanistic understanding of co-aggregation between oral bacteria, one of the key organization principles driving dental plaque formation.

## RESULTS

### *V. parvula* uses specific adhesins to interact with *S. oralis*, *S. gordonii* and *A. oris*

In order to identify potential ligands of *V. parvula* SKV38 adhesins, we used our model *V. parvula* SKV38 strain to perform co-aggregation assays with different bacterial members of the dental plaque. *V. parvula* SKV38 co-aggregated with several *Streptococcus gordonii* strains, *Streptococcus oralis* ATCC10557 and *Actinomyces oris* CIP102340. It did not, however, co-aggregate with *Streptococcus mitis* CIP 104996, *Streptococcus parasanguinis* CIP104372T, *Fusobacterium nucleatum* ATCC 25586 and *Streptococcus mutans* NG8, UA159, CBSm8 and CBSm38 and only very weakly with *S. mutans* UA140 (Figure 1A, Figure S1). We decided to further investigate the determinant of co-aggregation between *V. parvula* SKV38 and *S. oralis* ATCC 10557, *S. gordonii DL1* and *A. oris* CIP102340. To identify which of the 12 *V. parvula* adhesins were involved in the co-aggregation with these different partners, we used previously constructed single deletion mutants of each of these adhesins^25^ and performed co-aggregation assays by mixing independent cultures of each of the three tested oral bacterial strain and the 12 *V. parvula* adhesin mutants in aggregation buffer. Deletion of *V. parvula* trimeric autotransporter VtaA abolished co-aggregation with *S. oralis,* while deletion of the trimeric autotransporter VtaE abolished co-aggregation with *S. gordonii* and strongly reduced co-aggregation with *A. oris* (Figure 1B-E and S2). A double mutant lacking both VtaA and VtaE showed reduced co-aggregation with *A. oris* compared to a *ΔvtaE* single mutant, suggesting that VtaA is a secondary adhesin involved in the co-aggregation with *A. oris* (Figure 1G). Microscopy observation of *V. parvula* incubated with *S. oralis, S. gordonii* and *A. oris* confirmed the observed co-aggregation phenotypes (Figure S2). Moreover, use of P_Tet_-*vtaA* or P_Tet_-*vtaE* constructs, in which the chromosomal *vtaA* and *vtaE* genes are placed under the control of an aTc inducible promoter, allowed us to recapitulate the aggregative phenotype in an aTc-dependent manner (Figure 1C-E). Both the P_Tet_-*vtaA* and the P_Tet_- *vtaE* strains partially co-aggregated with *S. oralis* and *A. oris*, even in absence of aTc, suggesting a leakage of the used P_Tet_ promoter. While deletion of *vtaE* completely abolished co-aggregation with *S. gordonii* when mixed after independent growth, it only partially abrogated co-aggregation with *S. gordonii* when cocultured overnight (Figure 1F), suggesting that another *V. parvula* adhesin could contribute to co-aggregation. Consistently, we identified VtaD as being this secondary adhesin, since any residual co-aggregation between *S. gordonii* and *V. parvula* disappeared in the Δ*vtaCDEF* and Δ*vtaDE* mutants (Figure 1F). *vtaD* is the gene located immediately upstream of *vtaE* and VtaD has a high similarity to VtaE (81%), which may explain why both corresponding proteins possess similar binding activities. However, *vtaD* encodes a shorter adhesin than VtaE (2071 residues opposed to 3141 residues), mostly lacking part of the repetitive sequences found in *vtaE* stalk (Figure S3 and S4). Interestingly, deletions of *vtaC* or *vtaF* in the Δ*vtaE* background increased the aggregative phenotype of *V. parvula* with *S. gordonii* (Figure 1F) suggesting that these other adhesins may interfere with the VtaD-dependent co-aggregation process.

**Figure 1:**
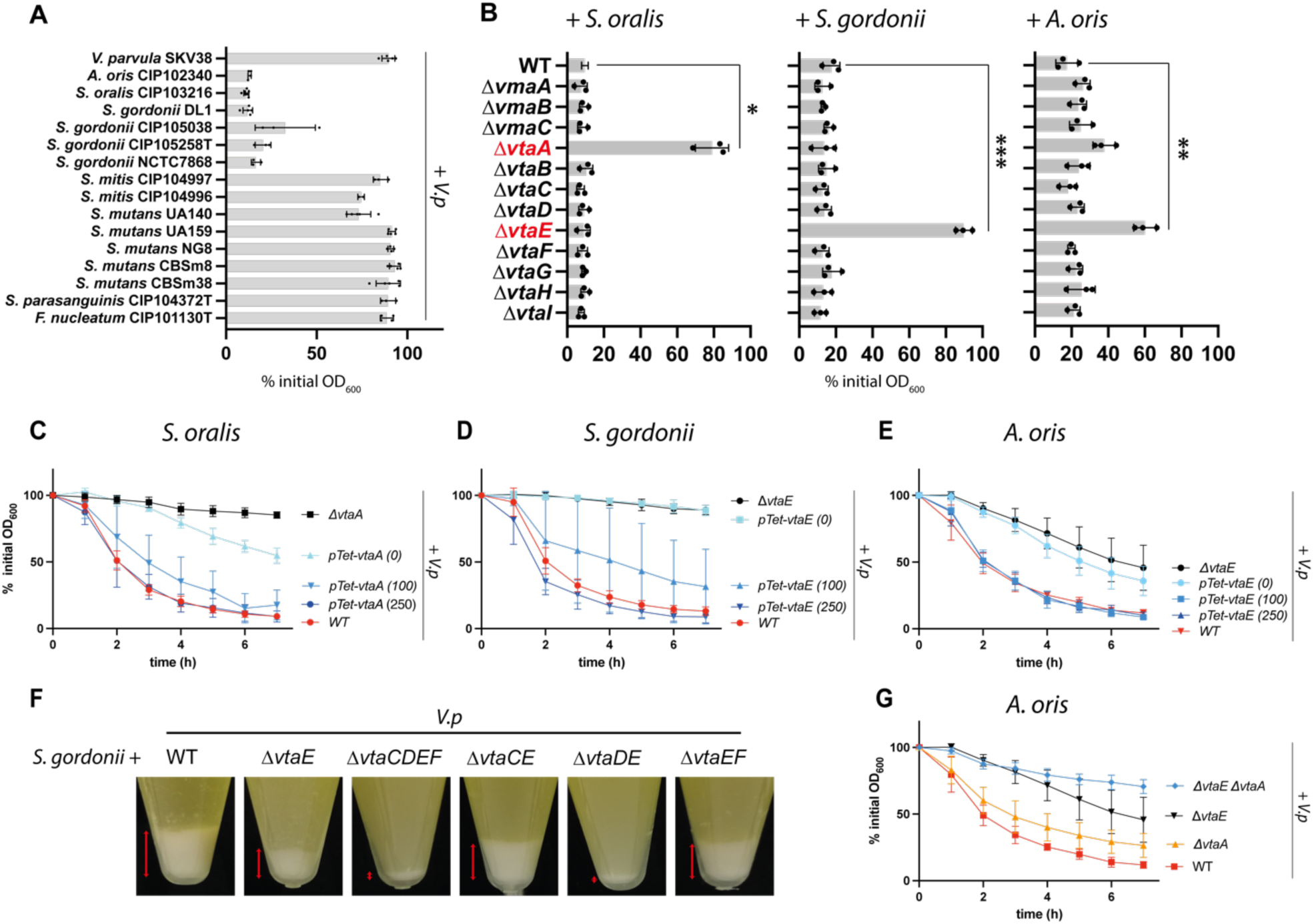
VtaA and VtaE are the adhesins responsible for co-aggregation with *S. oralis, S. gordonii and A. oris.* (A) Co-aggregation of independent cultures of both *V. parvula* SKV38 and various members of the dental plaque after 7h, as measured by the % of decrease of optical density between 0 and 7h. SD and single points for 3-5 replicates are shown. See Figure S1 for auto-aggregation of each strain. (B) Aggregation of *V. parvula* SKV38 WT and each single autotransporter mutant with *S. oralis ATCC 10557, S. gordonii* DL1 and *A. oris* CIP102340 after 7h. SD and single points for 3 replicates are shown. The indicated p-values were calculated by comparing all conditions to the partner + *Vp* WT using a Brown-Forsythe and Welch ANOVA followed by Dunnett correction. (C-E and G) Co-aggregation curves of *V. parvula* WT, Δ*vta*A, Δ*vtaE, ΔvtaEΔvtaA* and P_Tet_ -*vtaE* or P_Tet_ -*vtaA* with 0, 100 or 250 ng/ µl of aTc. Curves represent the mean and SD of 6-17 replicates. (F) Representative pictures of co-aggregates after coculture between *S. gordonii* WT and *V. parvula* WT and different adhesin mutants; red arrow bars indicate the relative size of the aggregated fraction.

The Hag1 trimeric autotransporter has been shown to be involved in the adhesion of *V. atypica* OK5 to human oral epithelial cells^23^. Interestingly, the genes encoding VtaA and Hag1 are located at the same locus on the genome of *V. parvula* SKV38 and *V. atypica* OK5, respectively, with the difference that Hag1 is preceded by another trimeric adhesin. Comparison of this locus among different *Veillonella* revealed that this locus always contains adhesins, although the number of adhesin and their identity differs between strains, even within the same species (Figure S5). This feature is reminiscent of *V. parvula* SKV38^25^ cluster of adhesin that is also present in a locus that consistently hosts diverse adhesins across *Veillonella* species.

Apart from its importance in the dental plaque, *V. parvula* is also present throughout the gastrointestinal tract. We wondered whether some of its adhesins are involved in adhesion to oral or intestinal cells, rather than other bacteria. In contrast to the known strong interaction between *V. atypica* and host cells^23^, we observed only a moderate adhesion of *V. parvula* SKV38 to TR146 oral and Caco-2 intestinal epithelial cells using microscopy (Figure S6 A-C). We then tested whether the major adhesins of *V. parvula* were involved in this interaction using a Δ*vtaCDEF*Δ*vtaA* mutant.. Deletion of the large adhesin group did not reduce adhesion to either cell type. Finally, we examined whether the other adhesins of *V. parvula* SKV38 could impact Caco-2 cell adhesion, and showed that there were no significant differences in adhesion (Figure S6D).

### Identification of VisA (*SGO_2004*), a new *S. gordonii* adhesin mediating co-aggregation with *V. parvula*

To further characterize the molecular actors of co-aggregation, we focused on the pair *V. parvula / S. gordonii* and took advantage of a recently published collection of 27 *S. gordonii* DL1 surface proteins deletion mutants^26^, corresponding to all 26 LPXTG cell wall anchor domain-containing proteins plus two mutants of the Amylase-binding protein A (AbpA) and B (AbpB). We first investigated co-aggregation between wild-type *V. parvula* and all *S. gordonii* mutants and we identified two mutants, Δ*padA* (*SGO_2005*) and Δ*SGO_2004*, presenting either a reduced (Δ*padA*) or total loss of co-aggregation (Δ*SGO_2004*) with *V. parvula* (Figure 2A-B).

**Figure 2:**
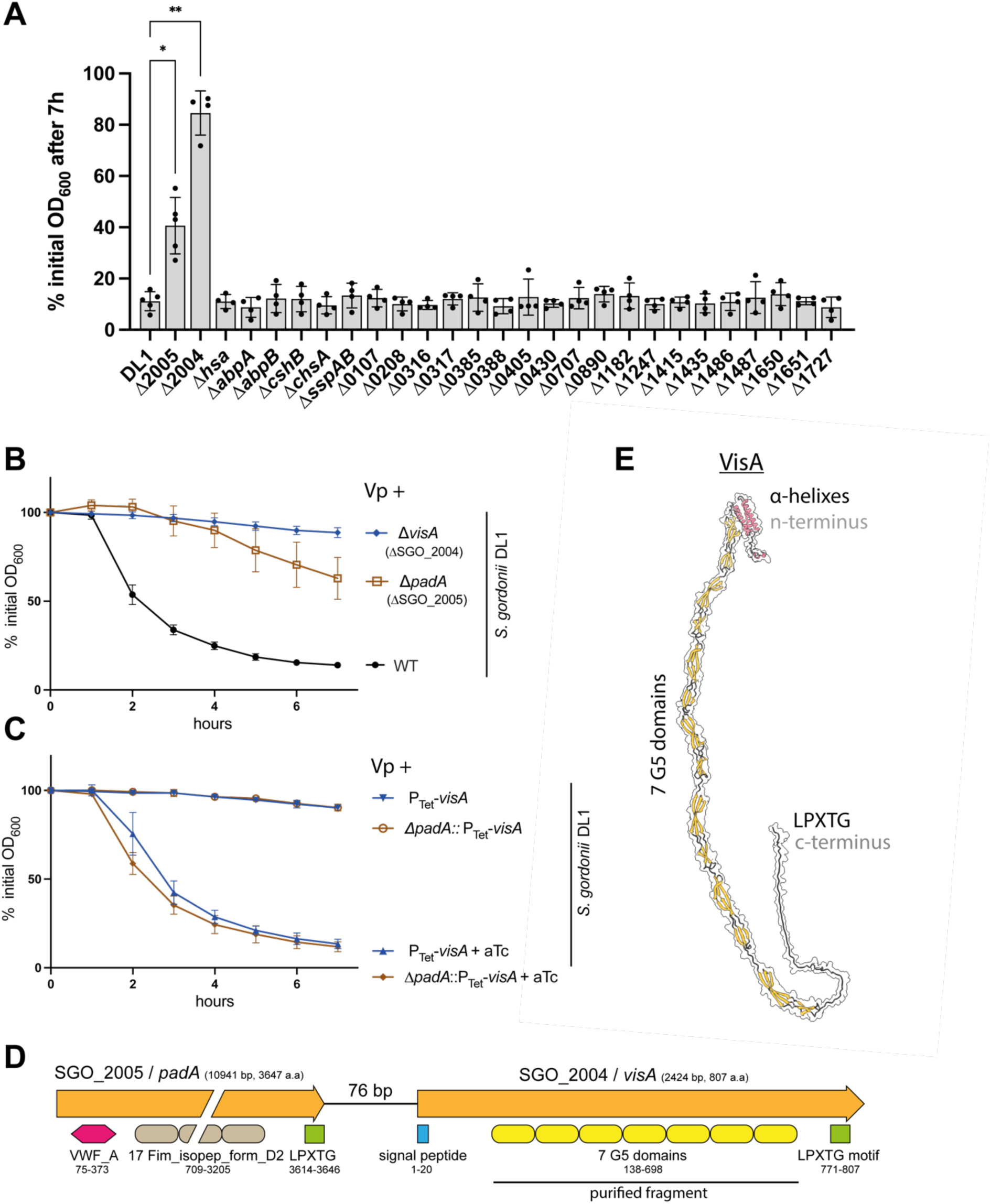
VisA (SGO_2004) is a novel adhesin interacting with *V. parvula.* (A) Co-aggregation of *V. parvula* SKV38 with *S. gordonii* DL1 WT and mutants for each LPXTG-containing protein and *abpA-B*, as measured by the % of decrease of optical density between 0 and 7h. SD and single points for 4-5 replicates are shown. The indicated p-values were calculated by comparing all conditions to the partner + *Vp* WT using a Brown-Forsythe and Welch ANOVA followed by Dunnett correction. Co-aggregation curves of *S. gordonii* WT, Δ*visA, ΔpadA* (B) and P_Tet_-*visA* or P_Tet_-*padA* (C) with or without 250 ng aTc. Curves represent the mean and SD of 6-13 replicates. (D) Genetic organization of the SGO_2004/2005 locus. VWF_A: Von Willbrand factor A (IPR002035), Fim_iso-pep_form_D2: Fimbrial isopeptide formation D2 domain (IPR026466), G5 domain (IPR011098). (E) AlphaFold structural model of VisA without the signal peptide.

*padA* and *SGO_2004* are part of an operon (Figure 2D) and the observed loss of aggregation in the *ΔpadA* mutant could be due to a polar effect on the downstream *SGO_2004* gene^27^. To test for this hypothesis, we inserted a P_Tet_ inducible promoter with the pVeg RBS ^28^ upstream of *SGO_2004,* while retaining or deleting the *padA* gene. In both cases, co-aggregation was fully recovered in presence of aTc (Figure 2C), demonstrating that SGO_2004 alone is the protein responsible for *S. gordonii* co-aggregation with *V. parvula*. *SGO_2004* is a gene of previously unknown function coding for an 807 amino acid protein composed of a flexible chain of disordered/poorly predicted 3 short alpha helixes, 7 G5-domains and an LPXTG domain (Figure 2D-E). Homologues of this protein are found in other, sometime distant, Streptococci, next to a *padA* homologue (Figure S7). Considering its newly identified role, we renamed this new aggregation-mediating adhesin VisA, for **V**eillonella **I**nteracting **S**treptococcal protein A.

### *S. gordonii* VisA directly interacts with *V. parvula* VtaE and VtaD

To determine whether co-aggregation mediated by *V. parvula* VtaE and VtaD and *S. gordonii* VisA resulted from direct or indirect interactions, we purified the VisA region containing its 7 G5 domains (residues 138-698 with a C-terminal His-tag, see Figure 2D) in *E. coli* and used the purified protein to assess potential direct interactions with *V. parvula*. When used at a concentration above 1µg/mL, VisA_G5_ was sufficient to induce aggregation of *V. parvula* on its own (Figure 3A). Confirming our previous observations, a Δ*vtaE* mutant retained a partial aggregation phenotype, while a Δ*vtaD-vtaE* mutant did not, and Δ*vtaE*Δ*vtaC* and Δ*vtaE*Δ*vtaF* mutants displayed an intermediate phenotype (Figure 3B). Moreover, immunofluorescence using an anti-His antibody detecting VisA_G5_ incubated with *V. parvula* WT, *ΔvtaE* or *ΔvtaD-vtaE* showed that while VisA_G5_ could be detected at the surface of *V. parvula* WT (Figure 3C) or *ΔvtaE* (Figure 3D), no signal could be seen for the *ΔvtaD-vtaE* mutant (Figure 3E). Altogether, these results suggested that VisA binds to *V. parvula* surface via a direct interaction with VtaE or VtaD.

**Figure 3:**
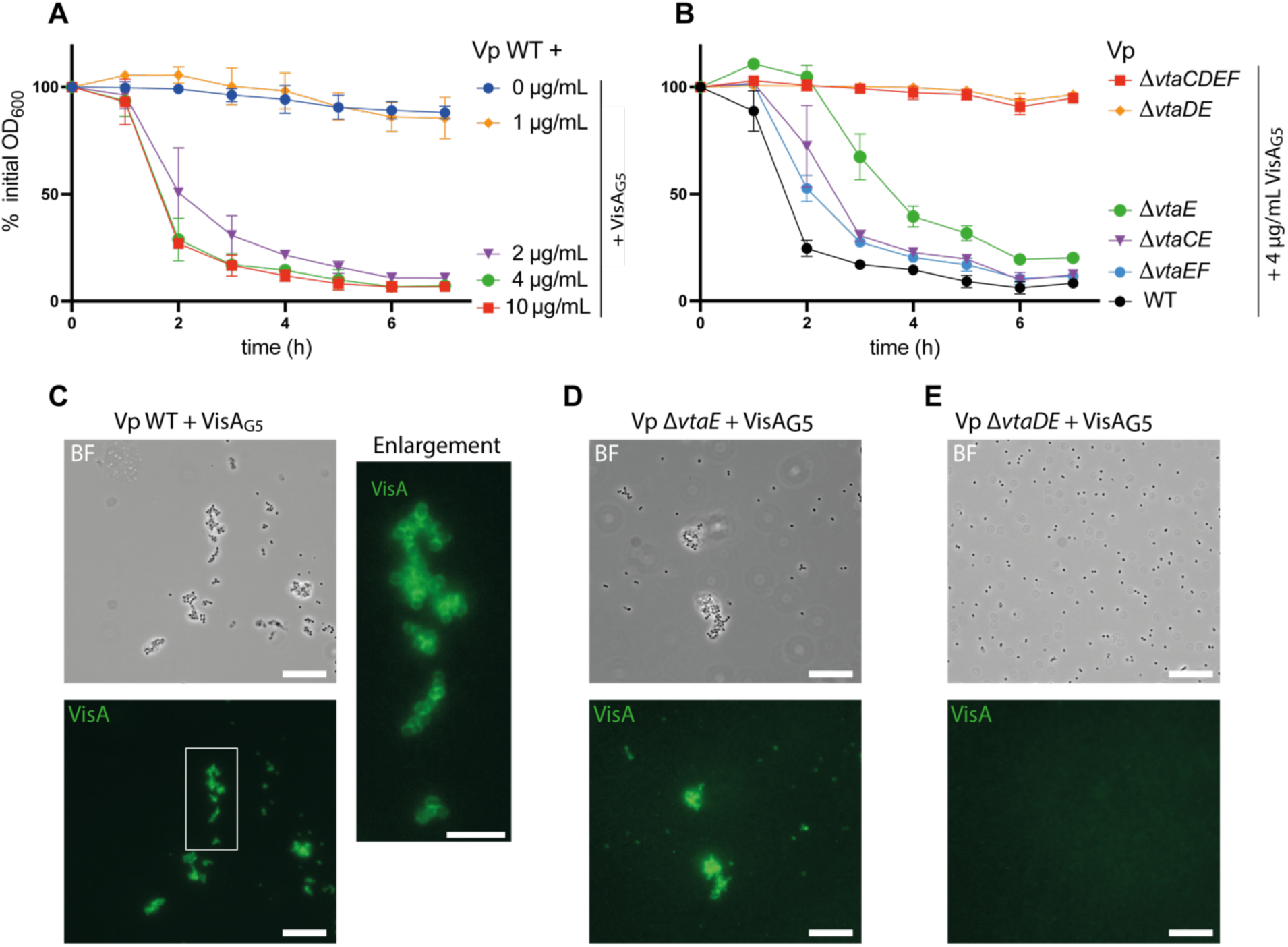
VisA binds directly to *V. parvula* by interacting with VtaE and VtaD. (A) Auto-aggregation curves of *V. parvula* SKV38 with various concentrations of VisA_G5_. (B) Aggregation curve of *V. parvula* SKV38 or indicated adhesin mutants with 4 µg/mL of VisA_G5_. For (A) and (B), curves represent the mean and SD of 3 replicates. (C-E) Brightfield images and their corresponding immunofluorescence images targeting the His-tag of VisA_G5_ after incubation of *Vp* WT, Δ*vtaE* and Δ*vtaDE* with 10 µg/mL of VisA_G5_ protein. Scale bar is 15 µm. The (C) right panel represents an enlargement of WT + VisA_G5_ immunofluorescence image (indicated by the white square) and scale bar is 5 µm.

### Co-aggregation in co-culture produces no significant alteration on the transcriptomic profiles of *V. parvula* and *S. gordonii*

While previous studies have compared the transcriptional responses of *Veillonella* and *S. gordonii* co-incubations compared to mono-incubation^18,19,29^, they did not specifically evaluate the potential contribution of co-aggregation. Having identified the adhesins involved in *V. parvula / S. gordonii* co-aggregation, we set out to compare the transcriptional responses of these two strains in mono- and cocultures with and without co-aggregation or auto-aggregation. Here we used the rich medium BHIP (BHI + 100 mM pyruvate), in which both bacteria could grow without metabolic co-dependency.

*V. p*a*rvula* transcriptional profiles of each condition grouped mainly by the presence of *S. gordonii* and then by their strain type. In principal component analysis (PCA), calculated using normalized transcripts counts, samples were strongly separated on the first principal component by their coculture status, thus indicating that the main determinant of the observed *V. parvula* response is the presence of its bacterial partner *S. gordonii* (Figure 4A). The PCA analysis on the second and third axis revealed a clustering by *V. parvula* mutant (Figure S8), suggesting that the residual differences between conditions are associated with the nature of the *V. p*a*rvula* mutants.

**Figure 4:**
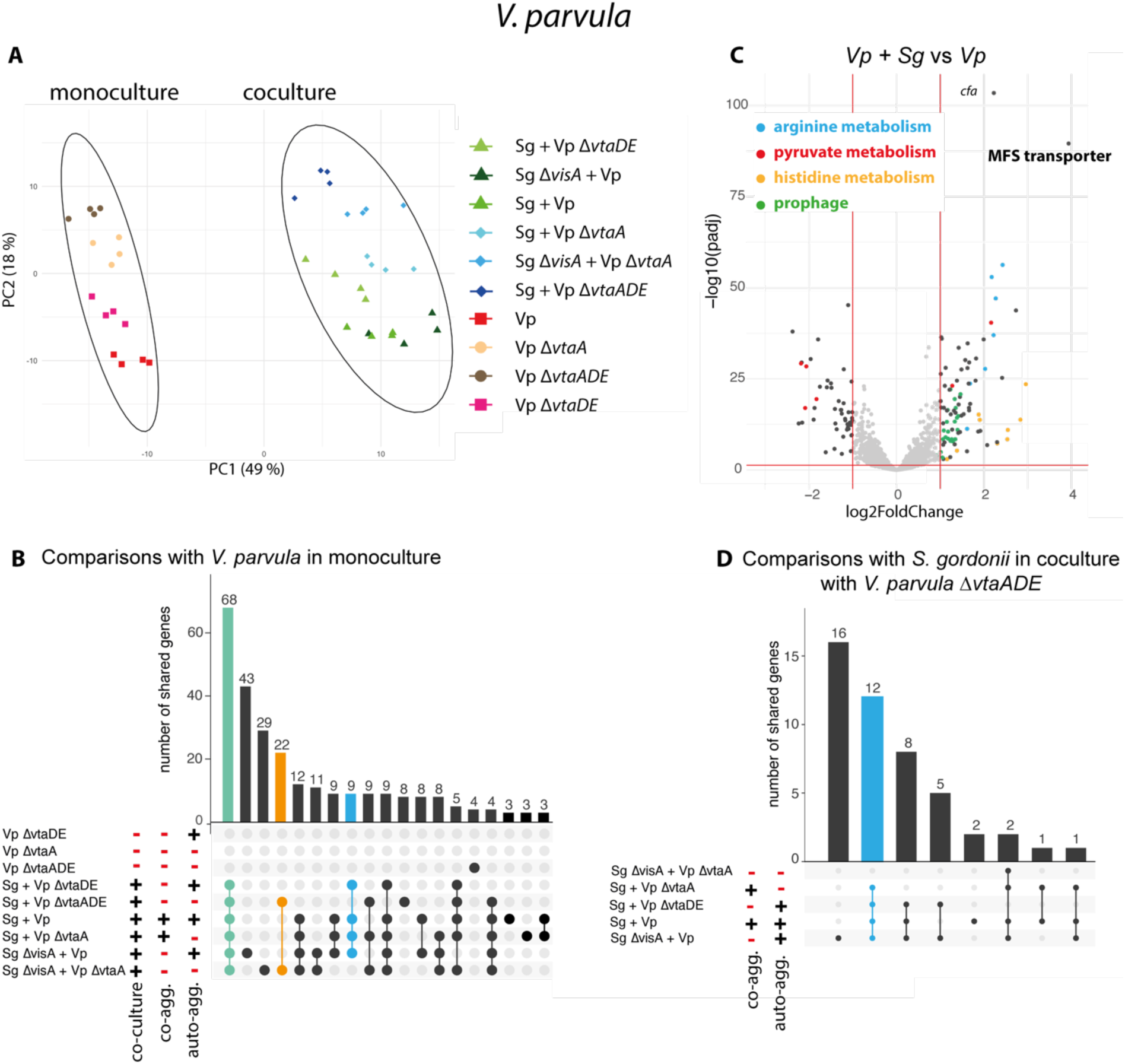
Transcriptomic response of *V. parvula* to *S. gordonii* is mostly related to coculture. A) Principal component analysis (PCA) of all *V. parvula* samples (4 biological replicates for 10 conditions). Colors and shape represent the different conditions. The two circles separate monoculture samples from coculture samples. Green symbols indicate samples able to auto-aggregate in coculture, blue shades samples unable to auto-aggregate. B) Upset plot (a Venn diagram alternative) showing the number of differentially expressed genes (defined by an absolute log2fold change > 1) shared for each condition compared to *V. parvula* WT monoculture. The green bar indicates the core response to coculture, the orange bar the core answer to coculture without any aggregation and the blue bar the response to any aggregation in coculture. C) Volcano plot of the coculture of *V. parvula* and *S. gordonii* WT compared to *V. parvula* in monoculture. Genes corresponding to identified key functions are differentially colored. D) Upset plot for each condition compared to *V. parvula* Δ*vtaAΔvtaDE* and *S. gordonii* coculture., the blue bar shows the response to any aggregation in coculture.

In order to identify potential coculture-specific response, we searched for genes up or downregulated (log2Fold above 1 or below −1) in at least one condition compared to *V. parvula* WT monocultures. The resulting Upset plot (Figure 4B) represents the common dysregulated genes for different combinations of conditions. This plot shows that the core *V. parvula* coculture transcriptomic response in all conditions was composed of 68 genes (Figure 4B green bar and supplementary data S1). The most upregulated gene was *FNLLGLLA_00352* (around 4.5 log2Fold increase compared to the monoculture), coding for an uncharacterized major facilitator superfamily-type (MFS) transporter, an inner membrane transporter of an unknown small molecule. We also found a strong upregulation of genes coding for enzymes of the histidine and arginine biosynthesis pathways (Figure 4C). Interestingly, *vtaB*, encoding an uncharacterized trimeric autotransporter and a gene cluster encoding a prophage were also induced, albeit at lower levels. Many genes associated with stress response were slightly upregulated (genes coding for the chaperones GroEL and GroES, their regulators CtsR and HcrA, ClpC and ClpE) (supplementary data S1). Pyruvate metabolism appeared to be remodeled in coculture by up- and downregulation of many pyruvate-associated genes (Figure 4C, supplementary data S1). Concerning lactate consumption, the malate/lactate antiporter *mleN* was slightly up-regulated, while genes related to the L- and D-lactate dehydrogenases were downregulated (*lutA*-*lutC*, *FNLLGLLA_01898* and *fucO*). Genes involved in iron or other metal uptake through the inner membrane were also both up- and downregulated.

We also compared specifically all coculture conditions compared to *V. parvula* Δ*vtaADE* with *S. gordonii* (Figure 4D). Overall, only a few *V. parvula* genes involved in purine metabolism were upregulated specifically wen aggregating in cocultures, either through co-aggregation or auto-aggregation (Figure 4B and D, blue bar, supplementary data S1). By contrast, 22 genes were specifically dysregulated in coculture in absence of any type of aggregation among which genes involved in NADH regeneration through xanthine to urate conversion were slightly downregulated (Figure 4B, orange bar, supplementary data S1).

On the other hand, there were very few changes on *S. gordonii* transcriptome when cocultured with *V. parvula*. The only upregulated genes in all cocultures conditions (Figure 5AB, green bar, supplementary data S2) are part of the Bfb PTS system (SGO_1575-82) already described as induced when co-aggregating with *A. naeslundii*^30^. The only downregulated gene (SGO_1314) encoded a ZnuA-like metal binding lipoprotein (Figure 5C). No gene expression changes were found specifically associated to co-aggregation (Figure 5D).

**Figure 5:**
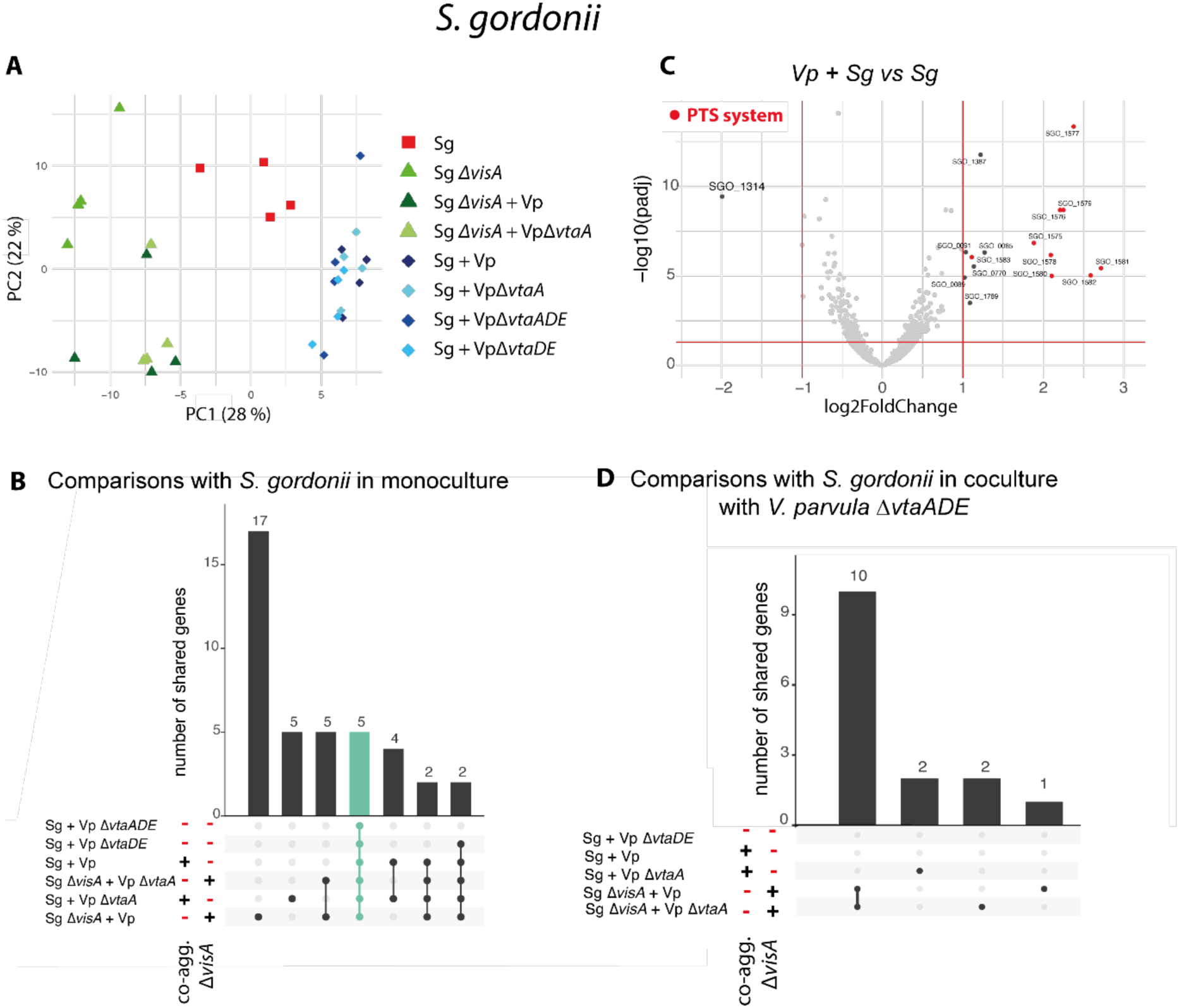
Transcriptomic response of *S. gordonii* to *V. parvula* is limited to the upregulation of a PTS system and downregulation of a metal binding lipoprotein. A) Principal component analysis (PCA) of all *S. gordonii* samples (4 biological replicates for 10 conditions). Shades of green represent all *S. gordonii* ΔvisA conditions, shades of blue cocultures of *S. gordonii* WT and red the monoculture of *S. gordonii* WT. B) Upset plot (Venn diagram alternative) showing the number of differentially expressed genes (defined by an absolute log2fold change > 1) shared for each condition compared to *S. gordonii* WT or *ΔvisA* monocultures (indicated by the *ΔvisA* column). The green bar indicates the core response to coculture and the blue bar the core differences between *S. gordonii* WT and Δ*visA*. C) Volcano plot of the coculture of *V. parvula* and *S. gordonii* WT compared to *S. gordonii* in monoculture. Genes of the PTS system are colored in red. D) Upset plot for each coculture condition compared to *S. gordonii + V. parvula ΔvtaAΔvtaDE* coculture.

Altogether, these results indicate that (i), *V. parvula* transcriptional response to coculture is associated with changes in metabolism and stress (ii) *S. gordonii* has a minimal transcriptional response, (iii). aggregation has only a limited effect on both bacteria, without contribution of auto- or co-aggregation.

### Co-aggregation strongly affects the structure of mixed *V. parvula*/*S. gordonii* biofilms

To assess the impact of co-aggregation on mixed biofilm formation, we imaged either mono-species or mixed biofilms of *V. parvula* and *S. gordonii* formed in 96 well plates for 24h using confocal laser scanning microscopy (CLSM). To differentiate both bacteria, *S. gordonii* was stained using the monoderm specific dye BacGO^31^ while Syto61 was used to stain all bacterial (Figure S9A-B). Comparison of co-aggregating mixed biofilms (*Vp* WT + *Sg* WT) with mixed biofilms without co-aggregation (*Vp* Δ*vtaDE* + *Sg* Δ*visA*, *Vp* Δ*vtaDE* + *Sg* WT and *Vp* WT +*Sg* Δ*visA*) showed that, in absence of co-aggregation, the two partner bacteria were found in distinct patches (Figure 6A). This was confirmed by the measurement of roughness (capturing the variations of height over the biofilm) of the streptococcus biofilm in mixed biofilms (Figure 6B, figure S9C-D). However, co-aggregating biofilms presented a more homogenous distribution of the two bacterial populations (Figure 6A). Volume measurements were variable but suggested that co-aggregation results in a higher overall biofilm volume and an increased *S. gordonii* biofilm (Figure S9E-F). Measures of total biofilm formation by crystal violet assay did not show an increase in biofilm formation when co-aggregating (Figure S9G). Coaggregation therefore seems to strongly impact on the organization of the two species in mixed biofilms which could profoundly modulate the behavior of these species *in vivo*.

**Figure 6:**
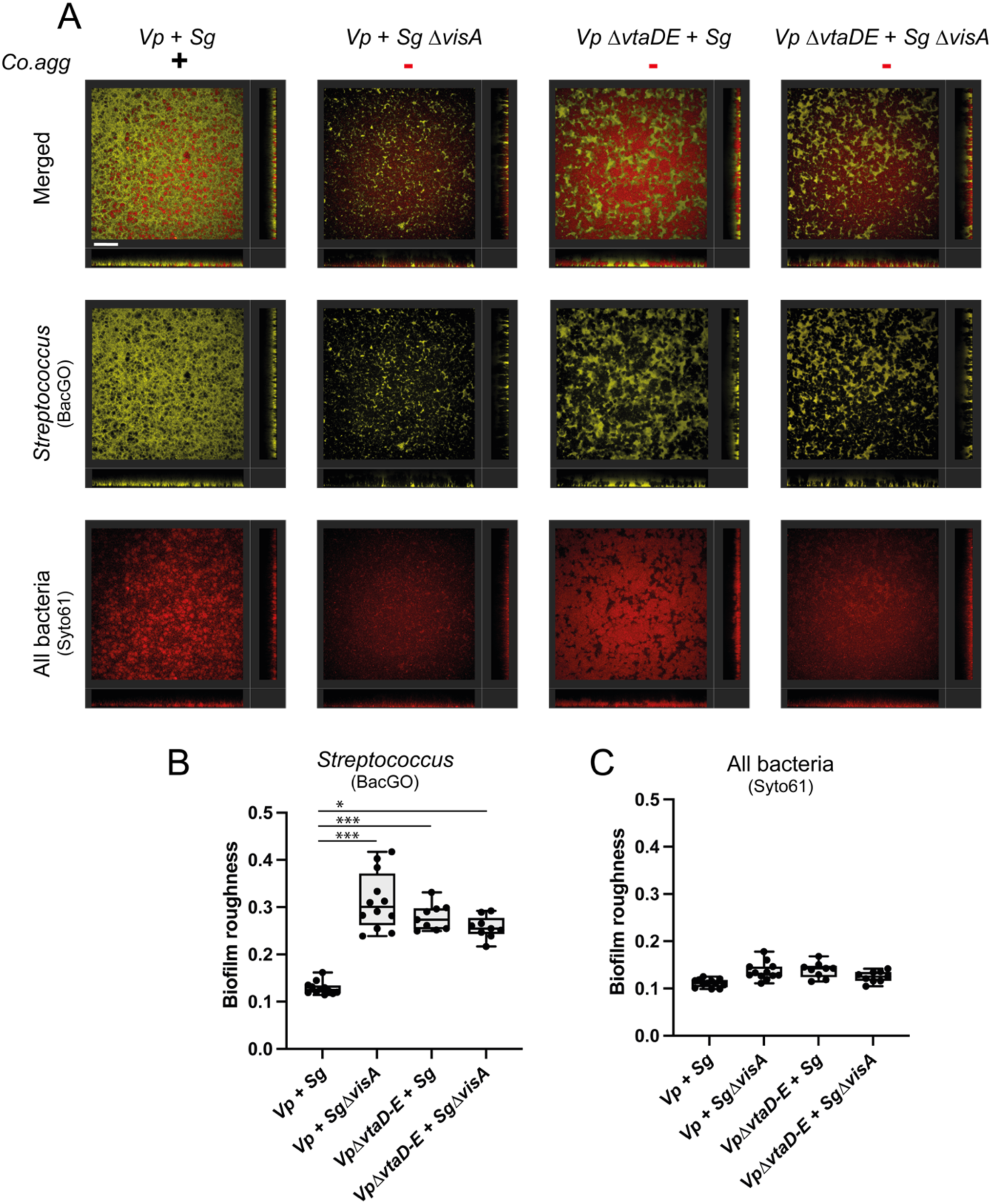
Confocal microscopy of mixed biofilms. A) Representative section images of mixed biofilms, scale bar is 60 μm. Lower and side bands correspond to the orthogonal projections on the z-x and z-y axis respectively. B-C) Measured biofilm roughness parameter for the BacGO and the Syto61 dyes. Each point (9-12 per condition) represents the average roughness measurement of four images per well. Experiment done in three biological independent replicates and three technical replicates. P-values, indicated by asterisk (*: p< 0,05, *** : p< 0,0005) were calculated using a Kruskal-Wallis test with Dunn’s correction for multiple testing. For all plots, *Vp* is *V. parvula*, *Δ*DE is *V. parvula* Δ*vtaD-vtaE*, *Sg* is *S. gordonii* and Δ*visA* is *S. gordonii ΔvisA*. Presence (or absence) of auto- and co-aggregation is indicated by the + (or −) symbols.

## DISCUSSION

Interactions between bacteria and their environment, whether abiotic or biotic, play a key role in determining the nature and evolution of bacterial lifestyles and we previously characterized the *V. parvula* adhesins involved in its biofilm formation capacities. In this study, we investigated the molecular determinants at the origin of the co-aggregation mechanisms between *V. parvula* and different members of the dental plaque and identified three *V. parvula* and one new *S. gordonii* adhesins involved in co-aggregation and studied the impact of such co-aggregation on partner physiology and co-biofilm structure.

### Adhesion strategies in *Veillonella*

We showed that the previously identified *V. parvula* VtaA adhesin interacts with *S. oralis* and *A. oris* while VtaE is responsible for co-aggregation with *A. oris* and *S. gordonii*, in which the highly homologous, but truncated VtaD has a secondary contribution (Figure 7).

**Figure 7:**
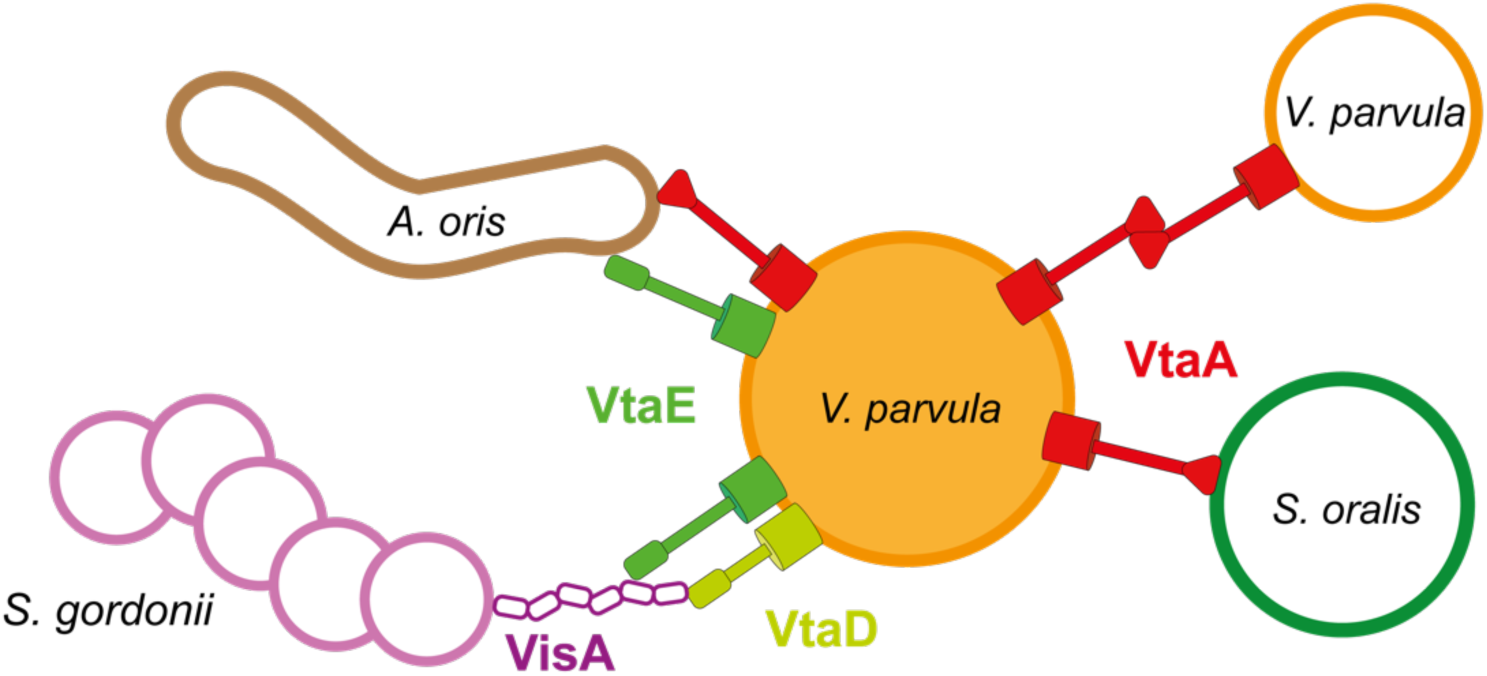
The multiple roles of *V. parvula* adhesins. Model of the interactions mediated by the different *V. parvula* adhesins.

Contrary to what has been described for *V. atypica*, where a single adhesin, Hag1, is responsible for all aggregative phenotypes^23^, the different adhesive functions in *V. parvula* are located on different proteins. Comparison of the predicted structures of Hag1 with VtaE, VtaD and VtaA shows that Hag1 head section is much more developed than the other adhesins, which could explain its pleiotropic role (Figure S3B). In addition, Hag1 is almost twice the size in residues (7187 residues) compared to Hag2 (3838 residues), the second longest adhesin in *V. atypica* OK5. In *V. parvula* SKV38, all major adhesins, including VtaA (3041 residues), VtaE (3142 residues), VtaC (2811 residues) and VtaF (3193 residues) are of similar size. One hypothesis is that Hag1, because of its long size, could mask other adhesins at the cell surface thus explaining the concentration of activities on the only surface accessible adhesin. By contrast, in *V. parvula* SKV38, activities are distributed across multiple co-expressed surface adhesins. VtaD and VtaE head domains are very similar, but VtaE is estimated to be around 100 nm longer than VtaD (Figure S3A) and we observed that more discrete aggregative phenotypes are associated with VtaD compared to VtaE, which could be due to masking of VtaD by VtaE. Additionally, the fact that the double mutants Δ*vtaE*Δ*vtaF* and Δ*vtaE*Δ*vtaC* aggregate faster with the purified *S. gordonii* VisA_G5_ than the simple Δ*vtaE* mutant is also in favor of the hypothesis that a shorter VtaD adhesin is partially masked by the longer VtaC and VtaF adhesins. This masking interference between adhesins has been commonly observed as a possible regulatory mechanism of the surface structures ^15–16^. Therefore, selection pressure on adhesion could either apply towards ensuring that the main adhesins do not mask each other by remaining of similar size, still allowing some potential interference relief of shorter adhesins (*V. parvula* case) or towards accumulating all functions on the tallest adhesin (*V. atypica* case).

Here we identified VtaA as an adhesin promoting co-aggregation with *S. oralis*, whereas we previously showed that VtaAit promotes auto-aggregation in BHI^34^. This auto-aggregation does not happen after growth in SK medium, which was used to grow *V. parvula* for co-aggregation assays. This switch from an auto-aggregative to a co-aggregative behavior depends on environmental conditions. This could be an efficient mean to rapidly adapt to abrupt changes of environment without affecting the quantity of a single adhesin at the cell surface.

Different *Veillonella* species occupy different niches within the oral microbiome. *V. parvula* is strongly associated with the dental plaque while *V. atypica* and *V. dispar* are found on soft surfaces. *Veillonella* HMT 780 has a strong specialization for keratinized gingiva^16^. It would be interesting to know if differential colonization sites stem from different co-aggregation capacities. This site specialization has been associated to certain genes (e.g., thiamine biosynthesis genes) but no difference in the number of adhesins between sites could be seen^16^. However, older studies have shown that *Veillonella* isolates from different origin within the mouth presented site specific co-aggregation capacities^7^. Revisiting the concept of strain-specific co-aggregation with a modern genetic approach leveraging genome sequencing and genetic manipulation, could help us decipher whether *Veillonella* adhesins specificity to different bacteria is related to the site specificity.

We found that *V. parvula* binds weakly to human epithelial cells. This differs to what has been described in *V. atypica*^23^. Therefore, different species of *Veillonella* might show different adhesion capacity to host cells, maybe linked to their isolation niche or their adhesin repertoire. We cannot discount that technical differences in our assay explain that difference. We used cancer cell lines, whereas Zhou *et al.* used buccal cells from a human buccal swab. In addition, the buccal cells were not thoroughly washed after adhesion as in our present protocol.

### VisA, a novel adhesin of *S. gordonii*

Like *V. parvula, S. gordonii* DL1 seems to use different adhesins to bind to different partners. For instance, it binds to certain *Veillonella* species, including *V. atypica* OK5, through Hsa, a sialic-acid binding protein also involved in platelet activation^24,32,33^. Here, we showed that Hsa is not involved in *S. gordonii* co-aggregation with *V. parvula* SKV38 (Figure 2A) and we have identified a second and new adhesin, VisA (SGO_2004), responsible for this interaction. Our results also suggest that VisA interacts directly with VtaE and VtaD. The use of purified VisA G5 domains demonstrated that they are the portion of VisA recognized by *V. parvula*. G5 domains are structural folds that are part of the stalk of monoderm surface proteins and are often found associated to an enzymatic active site^35^. For instance, SasG from *Staphylococcus aureus* or Aap from *Staphylococcus epidermidis* promote auto-aggregation through interaction between the G5-E domain repeats forming their B domain and have been described to undergo a zinc mediated dimerization^36^. While VisA does not seem to induce auto-aggregation of *S. gordonii*, the purified protein migrated exclusively at a size corresponding to a dimer in denaturing western blot (something also observed for trimeric autotransporters), suggesting that it also possesses the ability to dimerize even without the E-linker domain (Figure S10).

Interestingly, the locus encompassing genes encoding VisA-like proteins, PadA and a thioredoxin reductase is conserved in distant pathogenic *Streptococci* (Figure S7). PadA, in interaction with Hsa, is known to bind to platelets triggering their activation^33^. While in laboratory condition VisA (formerly known as SGO_2004) does not play a role in platelet interaction^27^, its conservation could suggest otherwise *in vivo*. *S. oralis* ATCC 10557 also possesses homologues of PadA (HRJ33_07090) and VisA (HRJ33_07095). However, *V. parvula* adhesins responsible for co-aggregation with the *S. gordonii* and *S. oralis* species are not the same, which strongly indicates that *S. oralis* likely uses a protein different from VisA to co-aggregate with *V. parvula*. The *S. oralis* VisA homologue possesses only five G5 domains while *S. gordonii* VisA has seven domains. The protein could be too short in *S. oralis* and masked by other surface components or not expressed. This could explain that VisA does not contribute to *S. oralis* co-aggregation with *V. parvula*.

Taken together, these results further illustrate the versality in the use of various adhesins to co-aggregate both for Streptococci and Veillonella species.

### What drives the response to coculture in oral bacteria?

Although limited, modifications of gene expression during coculture of *V. parvula* and *S. gordonii* were observed. For *S. gordonii*, the main answer to coculture with *V. parvula* was the upregulation of a PTS system encoded by the *bfb* operon (SGO_1575-1582). This system was found upregulated in *S. gordonii* when co-aggregating with *Actinomyces oris*^30^ and one gene of the operon downregulated when co-aggregating with *Fusobacterium nucleatum*^37^. The *bfb* operon is associated with biofilm formation as deletion of several genes led to a decrease in adhesion and biofilm formation while the operon promoter was 25% more active in biofilms^38^. An increase of arginine concentration could be at the origin of the induction of this *S. gordonii* PTS system. Indeed, arginine is known to be important for *S. gordonii* biofilm formation and arginine restrictions result in strong downregulation of the *bfb* operon in monoculture^39^. Co-aggregation of *S. gordonii* with *A. oris* resulted in downregulation of arginine biosynthesis and upregulation of the *bfb* operon through the uptake of *A. oris*-produced arginine. One of the upregulated pathways in *Veillonella* when cocultured with *S. gordonii* is arginine biosynthesis. Therefore, one can hypothesize that *V. parvula* would favor *S. gordonii* biofilm formation by producing arginine. We have, however, not detected any decrease in the arginine biosynthesis pathway in *S. gordonii* or changes in expression of arginine dependent regulators *argC, argR* or *argC*.

Globally, coculture did not result in major changes in gene expression in our experiments performed in anaerobic conditions using a rich and buffered media without metabolic dependency. Auto-aggregation and co-aggregation themselves had a negligible impact on the observed responses by both bacteria. The induction of the alpha-amylase *amyB* gene expression in *S. gordonii* caused by a an unknown diffusible signal produced by *V. parvula*^18,19^ was not observed in our experiments. This may be due to our specific conditions that did not allow production of the signal by *V. parvula*. Other examples of oral bacteria responding weakly to co-aggregation are *F. nucleatum* interacting with *S. gordonii*^37^ and *S. mutans* interacting with *V. parvula*^40,41^. These results suggest that oral bacteria do not actually sense attachment to other bacteria but rather changes in nutrient availability and environment conditions such as pH or oxidative stress. Auto-aggregation and biofilm lifestyle is known to induce large metabolic changes in common aerobic bacteria, inducing genes involved in stress response and anaerobic metabolism in *E. coli*^42^ which seem mostly driven by oxygen gradients, as shown in aggregates of *P. aeruginosa* ^43^.

While anaerobic conditions could explain the limited response caused during coculture and interactions between *V. parvula* and *S. gordonii*, the exposure to oxygen could strongly impact the response to co-aggregation of anaerobic bacteria. Indeed, in another study looking at *S. gordonii and V. parvula* co-transcriptomes, Mutha *et al.* reported broad changes in *Veillonella* including a predominant response to oxidative stress with 39 out of 272 regulated genes associated with it while *S. gordonii* samples presented high inter-variability^29^. No common gene regulation could be detected between our results and their results, possibly due to different experimental settings, as they looked at response from short (30 min) aerobic co-aggregation in saliva while we looked at transcriptional responses after 6 h of anaerobic coculture. The aerobic conditions used during this short co-aggregation period could explain the strong *V. parvula* response to oxidative stress exacerbated by *S. gordonii*.

### Could the proximity within the biofilm enhance synergistic or antagonistic interactions?

We hypothesized that co-aggregation could influence localization of the two bacteria within the biofilm. Indeed, co-aggregation was necessary to promote colocalization of the two bacteria. Proximity within the biofilm would be essential to *Veillonella parvula* as it can favor the uptake of lactate by bringing it closer to the producer Streptococci, which would also protect it from oxidative stress by consuming the O_2_ locally. It could also favor signal transduction as demonstrated for the distance dependent induction of *S. gordonii amyB*.

Without co-aggregation, both bacteria were distant from each other in the biofilm. This could be explained by a passive clonal development but also by an active prevention of biofilm colonization by non-aggregating partners. This could have a strong effect *in vivo* by limiting the entry of non-co-aggregating members (including *S. mutans*) into the dental plaque biofilm while permitting the presence of cooperative partners in close vicinity. A similar mechanism has been demonstrated in *Vibrio cholerae*, where deletion of *rbmA*, the gene encoding RbmA, a matrix protein involved in mother-daughter cell cohesion, resulted in higher penetration by invaders as cells were less tightly packed in the biofilm^44^. Additionally, mixed biofilms between RbmA producers and deficient strains resulted in patchy structures reminiscent of our observation.

Mixed biofilms have often been described to increase stress resistance compared to single species biofilms. For instance, synergistic biofilm formation by four marine bacteria promoted protection to invasion by the pathogen *Pseudoalteromonas tunicata* and increased resistance to hydrogen peroxide and tetracycline compared to monospecies biofilms^45^. The resistance in a three-species biofilm was due to protective capacity of one of the resident members^46^. We hypothesize that, while mixed biofilms are already more stress-resistant, co-aggregation between members could further increase stress resistance.

In conclusion, we have shown that *V. parvula* uses specific sets of multiple trimeric autotransporters to specifically interact with other members of the oral dental plaque. While these adhesive capacities are not necessary for intercellular communication, they reduce distance between members of the biofilm. The co- aggregation phenomena are likely to contribute to the highly organized process of dental plaque formation by modulating the successive addition of interacting bacterial species.

## MATERIAL AND METHODS

### Growth conditions

Bacterial strains are listed in TABLE S1. *Streptococcus* spp. and *A. oris* were grown in brain heart infusion (BHI) medium (Bacto brain heart infusion; Difco*). V. parvula* was grown in BHI supplemented with 0.6% sodium dl-lactate (BHIL) or SK medium (10 g liter−1 tryptone [Difco], 10 g liter−1 yeast extract [Difco], 0.4 g liter−1 disodium phosphate, 2 g liter−1 sodium chloride, and 10 ml liter−1 60% [wt/vol] sodium dl-lactate; described in Knapp et al.^47^), in which it does not auto-aggregate. Bacteria were incubated at 37°C under anaerobic conditions in anaerobic bags (GENbag anaero; bioMérieux no. 45534) or in a C400M Ruskinn anaerobic-microaerophilic station. *Escherichia coli* was grown in lysogeny broth (LB) (Corning) medium under aerobic conditions at 37°C. When needed, 20 mg/L chloramphenicol (Cm), 200 mg/L erythromycin (Ery), 300 mg kanamycin (Kan) or 2.5 mg/L tetracycline (Tet) was added to *V. parvula* cultures, 5 mg/L Ery was added to *S. gordonii* cultures and 25 mg/L Cm or 100 mg/L ampicilin (Amp) was added to *E. coli* cultures. All chemicals were purchased from Sigma-Aldrich unless stated otherwise.

### *Veillonella parvula* natural transformation

From plate, cells were resuspended in 1 mL SK medium adjusted to an optical density at 600 nm (OD_600_) of 0.4 to 0.8, and 15 μL was spotted on SK agar petri dishes. On each drop, 1-5 μL (75 to 200 ng) linear double-stranded DNA PCR product was added. The plates were then incubated anaerobically for 24-48 h. The biomass was resuspended in 500 μL SK medium, plated on SK agar supplemented with the corresponding antibiotic, and incubated for another 48 h. Colonies were streaked on fresh selective plates, and the correct integration of the construct was confirmed by PCR and sequencing.

### *Veillonella parvula* mutagenesis and complementation

*V. parvula* site directed mutagenesis was performed as described by Knapp and al^47^ and Béchon et al^25^. Briefly, upstream and downstream homology regions of the target sequence and the *V. atypica* kanamycin (*aphA3* derived from the pTCV-erm^48^ plasmid under the *V. parvula* PK1910 *gyrA* promoter) or tetracycline resistance cassette were PCR amplified with overlapping primers using Phusion Flash high-fidelity PCR master mix (Thermo Scientific, F548). PCR products were used as templates in a second PCR round using only the external primers, resulting in a linear dsDNA with the antibiotic resistance cassette flanked by the upstream and downstream sequences. *vtaE* chromosomal complementation was done by inserting in the promoter region the previously described *Veillonella* P_Tet_ promoter^25^ associated with an erythromycin resistance cassette. Primers used in this study are listed in Table S2 in the supplemental material.

### Streptococcus gordonii natural transformation

25 µL of an O/N culture, 100 µL of heat inactivated horse serum (Sigma), 900 µL of THY Broth, 2 µL of competence specific peptide (1 mg/mL, DLRGVPNPWGWIFGR, synthetized by GenScript) and 1-5 µL of linear double-stranded DNA PCR product were mixed in a microcentrifuge tube, incubated anaerobically for 5 to 8 hours at 37°C and plated on selective agar medium for 1 to 3 days. Colonies were streaked on fresh selective plates, and the correct integration of the construct was confirmed by PCR and sequencing.

### Streptococcus gordonii complementation

In order to create a markerless mutant of *SGO_2004* with a P_Tet_ promoter, we took advantage of the described IDFC2 cassette^49^, containing an erythromycin resistance and a mutant *pheS* gene encoding the A314G missense mutation providing sensitivity to *p*-chlorophenylalanine (4-CP). Briefly, the IFDC2 cassette and homology regions before and after the promoter of *SGO_2004* was amplified from an *S. gordonii* strain containing IDFC2. PCR products were used as templates in a second PCR round using only the external primers, which generated a linear dsDNA with the IFDC2 cassette flanked by the upstream and downstream sequences. *Streptococcus gordonii* DL1 WT was transformed with this construct and selected for insertion of the cassette with erythromycin.

In a second time, the IDFC2 cassette was replaced by the P_Tet_ promoter of pRPF185 plasmid fused with the pVeg RBS^50^ by creating a construct with similar homologies regions than for the IFCD2 cassette or by using an homology region upstream of *padA* to create the Δ*padA,pTet*-*SGO_2004* mutant. After transformation of *S. gordonii* IDFC2-DL1 with either construct, counter selection was done on BHI + *p-*Cl-Phe plates and selected mutants verified by sanger sequencing and for sensibility to erythromycin.

### Aggregation assay

Overnight cultures were centrifuged for 5 min, 5000 g and resuspended in aggregation buffer^23^ (1 mM Tris-HCl buffer, pH 8.0, 0.1 mM CaCl_2_, 0.1 mM MgCl_2_, 150 mM NaCl) to a final OD_600_ of 1. 400 μL of each culture for co-aggregation or 800 μL for auto-aggregation were added to a microspectrophotometer cuvette (Fisherbrand) and left to sediment on the bench in the presence of oxygen, so no growth should occur. The OD_600_ was measured every hour in a single point of the cuvette using a SmartSpec spectrophotometer (Bio-Rad). OD_600_ were then normalized to the initial OD_600_ by the formula.

### Purification of *SGO_2004* G5 domains

The portion of *SGO_2004* coding for G5 domains (residues 138-698) was amplified from *S. gordonii* and the pET22b-HIS vector was linearized by PCR. The PCR products were then purified and annealed by Gibson reaction. The plasmid was dialyzed and transformed in electrocompetent *E. coli* DH5-alpha. After verification of the construct by sequencing, the plasmid was purified and transformed in *E. coli* BL21(DE3)-pDIA17. After growth to OD_600_ 0.4, cells were induced with 0.1 mM IPTG and grown for 3h at 37°C before harvesting. Cell pellet was frozen O/N then resuspended in Buffer A (30 mM Tris-HCl pH 7.5, 300 mM NaCl, 30 mM Imidazole) and lysed by sonication. Debris were pelleted by ultracentrifugation (50 000 g, 30 min) and supernatant run through a HisTrap 5 mL column on an AKTA Explorer (GE) against a gradient of imidazole (30-300 mM). The purified protein was assessed for purity by SDS-Page followed by SafeStain SimplyBlue^TM^ (ThermoFisher) staining and western blot against the HIS-tag (Figure S10) and dialyzed twice against 30 mM Tris-HCl pH 7.5, 300 mM NaCl using a SnakeSkin™ 3500 Da (ThermoFisher).

### Immunofluorescence of surface bound VisA_G5_

*V. parvula* was grown overnight in SK and washed two times in PBS. VisA_G5_ was preincubated 1 hour in the dark at 0.1 mg/mL with 1/10 of an anti-His Tag monoclonal antibody coupled with Alexa Fluor^TM^ 488 (MA1-135-A488, Invitrogen). 50 µL of bacteria at OD_600_ 1 was incubated for 2 hours with 5 µL of the fluorescent VisA_G5_ and subsequently mounted on a slide. Cells were imaged using a Zeiss Axioplan 2 microscope equipped with an Axiocam 503 mono camera (Carl Zeiss, Germany). Epifluorescence images were acquired using the ZEN lite software (Carl Zeiss, Germany) and processed using Fiji (ImageJ).

### RNA extraction

600 µL of anaerobic media BHIP (BHI + 100 mM sodium pyruvate) in a 1.5 mL tube was inoculated with each of the bacteria at OD_600_ 0.05 and incubated for 6h anaerobically. Resulting culture was mixed with 1.2 mL of RNAprotect Bacteria reagent (QIAGEN), vortexed and incubated at RT for 5 min, before centrifugation (10 000 rpm, 4°C) for 5 min. Supernatant was removed and pellet kept at −80°C before RNA extraction. For lysis, pellets were washed with 700 µL of PBS and resuspended in 200 µL of lysis buffer (15 mg/mL Lysozyme, 100 µL / mL Proteinase K) before incubation for 3h at 37°C with constant shaking (750 rpm). Each sample was then added to a matrix B lysis tube with 800 µL of TRIzol and lysed using a FastPrep (2 times *S. mutans* pre-registered protocol). 800 µL of 100% ethanol was added and samples were centrifuged to pellet debris (8000 g, 2min). Lysate was transferred to a column from the kit Direct-zol RNA Miniprep plus (Zymol) and the rest of the extraction was done following the providers manual.

### RNA sequencing

Libraries were prepared using Illumina Stranded Total RNA Prep from 440 ng of RNA. RiboZero kit Microbiome kit (Illumina) was used to eliminate ribosomal RNA. The subsequent steps were as follows: RNA fragmentation, cDNA synthesis (incorporating uracils into the second strand), adapter ligation, indexing by PCR with 17 cycles (amplifying only the first strand), purification of unbound adaptors and primers on AMP beads (Beckman Coulter). The resulting stranded libraries comprised fragments from 200 to 1000 bp with peaks lying between 390 and 470 bp as visualized on a 5300 Fragment Analyzer (Agilent Technologies). No low-molecular peaks corresponding to unbound adaptors and primer dimers were observed. Libraries were pooled and sequenced on a NovaSeq X 10 B flow cell (Illumina) producing 1200 millions 150×150-bp pair-end reads. As a result, each sample was represented by 18-55 million reads. Ribofinder was used to verify the efficiency of ribodepletion: only around 5% of reads mapped to ribosomal RNA. Taxonomy analysis using Kraken module confirmed the presence of *S. gordonii* and *V. parvula* RNA according to the co-infection design. In coinfection samples, reads from the two species were present in more or less equal proportions. The RNA-seq analysis was performed with Sequana^51^. In particular, we used the RNA-seq pipeline (v0.19.2, (https://github.com/sequana/sequana_rnaseq)) built on top of Snakemake v7.32.4^52^. Reads were trimmed from adapters and low-quality bases using fastp software v0.22.0^53^, then mapped to the reference genome using Bowtie2 v2.4.6^54^. Genomes and annotations were downloaded from NCBI website using *Veillonella parvula* SK38 (GenBank LR778174.1) and *S. gordonii* DL1 (GenBank CP000725.1) genome references. FeatureCounts 2.0.1^55^ was used to produce the count matrix, assigning reads to features using annotation aforementioned. Statistical analysis on the normalized count matrix was performed to identify differentially regulated genes. Differential expression testing was conducted using DESeq2 library 1.34.0^56^ scripts, and HTML reporting was made with the Sequana RNA-seq pipeline.Parameters of the statistical analysis included the significance (Benjamini-Hochberg adjusted p-values, false discovery rate FDR < 0.05) and the effect size (fold-change) for each comparison.

### Confocal Laser Scanning Microscopy

Biofilms were formed in a 96 well plate (PhenoPlate, PerkinElmer) by inoculating 150 µL of anaerobic media BHIP (BHI + 100 mM sodium pyruvate) with overnight culture of each species at OD_600_ 0.05 for each of them. After one hour of adhesion, media was replaced to remove planktonic bacteria and incubated for 24 hours. Biofilm was stained by addition of 50 µL of BHIP media containing both the BacGO (1 µM final concentration) and the Syto61 dies (5 µM final concentration). Three images set at defined positions within each well were acquired on an Opera Phenix Plus High Content Screening System running with Harmony software v.5.1 (Revvity, formerly known as PerkinElmer), using the following modalities: 20x water/NA 1.0, Z-stack, 40 planes, 2 µm step between planes, for the Syto61 dye: λ_exc_: 640 nm / emission filter 650-760 nm), for the bacGO dye: λ_exc_: 561 nm / emission filter 571-596 nm. Resulting images were analyzed using BiofilmQ 1.0.1^57^. Images were first denoised by convolution (dxy = 5, dz = 3) and top hat filter (dxy= 25), then segmented in two classes using an OTSU thresholding method with a sensitivity of 0.15 for the Syto61 channel and 0.25 for the BacGO channel. Images were then declumped in 10-pixel wide cubes and Surface properties (range 30 pixel) and Global biofilm properties calculated (supplementary data S3). Illustrative images were generated with Imaris 9.0.

## Data availability

Supplementary data are available at https://github.com/ldorison/Coaggregation_streptococcus_Veillonella-

## Acknowledgments

We thank Mark Herzberg for sharing the bank of *S. gordonii* surface proteins mutants, Justin Merritt for providing the *S. mutans* strains and the *S. gordonii* IDFC2 cassette, Robert Smith for his help with using the AKTA and developing the *Veillonella* kanamycin cassette and Bianca Audrain for developing the *Veillonella* erythromycin P_Tet_-cassette. We gratefully acknowledge the UTechS Photonic BioImaging (Imagopole), C2RT, Institut Pasteur, supported by the French National Research Agency (France BioImaging, ANR-10-INBS-04; Investments for the Future), and acknowledge support from Institut Pasteur for the use of the Revvity Opera Phenix Plus microscope.

This work was supported by Institut Pasteur and grants by the French government’s Investissement d’Avenir Program, Laboratoire d’Excellence “Integrative Biology of Emerging Infectious Diseases” (grant n°ANR-10-LABX-62-IBEID). L.D. was supported by a MENESR (Ministère Français de l’Education Nationale, de l’Enseignement Supérieur et de la Recherche) fellowship. C.M.G. was supported by a FRM Retour en France fellowship. The RNA sequencing and analysis was performed by the Biomics Platform, C2RT, Institut Pasteur, Paris, France, supported by France Génomique (ANR-10-INBS-09-09).

## Contribution

L.D., C.M.G. and C.B. designed the experiments. L.D., S.C., C.M.G., N.B., R.V., Y.V. and R.O. performed the experiments. L.D. and C.B. wrote the paper, with contributions from C.M.G., J.-M.G., N.B., R.O. and Y. V. and S.G. All authors read and approved the manuscript.

## Notes

https://github.com/ldorison/Coaggregation_streptococcus_Veillonella-

### Competing Interest Statement

The authors have declared no competing interest.

## Bibliography

1. Kolenbrander, P. E., Palmer, R. J., Periasamy, S. & Jakubovics, N. S. Oral multispecies biofilm development and the key role of cell–cell distance. Nature Reviews Microbiology 8, 471–480 (2010).

2. Kolenbrander, P. E. et al. Bacterial interactions and successions during plaque development. Periodontol 2000 42, 47–79 (2006).

3. Kolenbrander, P. E., Ganeshkumar, N., Cassels, F. J. & Hughes, C. V. Coaggregation: specific adherence among human oral plaque bacteria. The FASEB Journal 7, 406–413 (1993).

4. Afonso, A. C. et al. Bacterial coaggregation in aquatic systems. Water Research 196, 117037 (2021).

5. Hajishengallis, G., Lamont, R. J. & Koo, H. Oral polymicrobial communities: Assembly, function, and impact on diseases. Cell Host & Microbe (2023) doi:10.1016/j.chom.2023.02.009.

6. Mohanty, R., et al. Red complex: Polymicrobial conglomerate in oral flora: A review. J Family Med Prim Care 8, 3480–3486 (2019).

7. Hughes, C. V., Kolenbrander, P. E., Andersen, R. N. & Moore, L. V. Coaggregation properties of human oral Veillonella spp.: relationship to colonization site and oral ecology. Applied and Environmental Microbiology (1988) doi:10.1128/aem.54.8.1957-1963.1988.

8. Zhou, P., Manoil, D., Belibasakis, G. N. & Kotsakis, G. A. Veillonellae: Beyond Bridging Species in Oral Biofilm Ecology. Frontiers in Oral Health 2, (2021).

9. Kaplan, C. W., Lux, R., Haake, S. K. & Shi, W. The Fusobacterium nucleatum outer membrane protein RadD is an arginine-inhibitable adhesin required for inter-species adherence and the structured architecture of multispecies biofilm. Mol. Microbiol. 71, 35–47 (2009).

10. Takemoto, T. et al. Characteristics of multimodal co-aggregation between Fusobacterium nucleatum and streptococci. Journal of Periodontal Research 30, 252–257 (1995).

11. Guo, L., Shokeen, B., He, X., Shi, W. & Lux, R. Streptococcus mutans SpaP Binds to RadD of Fusobacterium nucleatum ssp polymorphum. Mol Oral Microbiol 32, 355–364 (2017).

12. Periasamy, S. & Kolenbrander, P. E. Central Role of the Early Colonizer Veillonella sp. in Establishing Multispecies Biofilm Communities with Initial, Middle, and Late Colonizers of Enamel. Journal of Bacteriology 192, 2965–2972 (2010).

13. Chung, W. O., Demuth, D. R. & Lamont, R. J. Identification of a Porphyromonas gingivalis receptor for the Streptococcus gordonii SspB protein. Infect Immun 68, 6758–6762 (2000).

14. Maeda, K. et al. Glyceraldehyde-3-Phosphate Dehydrogenase of Streptococcus oralis Functions as a Coadhesin for Porphyromonas gingivalis Major Fimbriae. Infection and Immunity 72, 1341–1348 (2004).

15. Coppenhagen-Glazer, S. et al. Fap2 of Fusobacterium nucleatum Is a Galactose-Inhibitable Adhesin Involved in Coaggregation, Cell Adhesion, and Preterm Birth. Infection and Immunity 83, 1104–1113 (2015).

16. Giacomini, J. J., Torres-Morales, J., Dewhirst, F. E., Borisy, G. G. & Mark Welch, J. L. Site Specialization of Human Oral Veillonella Species. Microbiol Spectr e0404222 (2023) doi:10.1128/spectrum.04042-22.

17. Delwiche, E. A., Pestka, J. J. & Tortorello, M. L. THE VEILLONELLAE: GRAM-NEGATIVE COCCI WITH A UNIQUE PHYSIOLOGY. Annual Review of Microbiology 39, 175–193 (1985).

18. Egland, P. G., Palmer, R. J. & Kolenbrander, P. E. Interspecies communication in Streptococcus gordonii–Veillonella atypica biofilms: Signaling in flow conditions requires juxtaposition. PNAS 101, 16917–16922 (2004).

19. Johnson, B. P. et al. Interspecies Signaling between Veillonella atypica and Streptococcus gordonii Requires the Transcription Factor CcpA. J Bacteriol 191, 5563–5565 (2009).

20. Zhou, P., Li, X., Huang, I.-H. & Qi, F. Veillonella Catalase Protects the Growth of Fusobacterium nucleatum in Microaerophilic and Streptococcus gordonii-Resident Environments. Applied and Environmental Microbiology 83, e01079–17 (2017).

21. Hughes, C. V., Andersen, R. N. & Kolenbrander, P. E. Characterization of Veillonella atypica PK1910 adhesin-mediated coaggregation with oral Streptococcus spp. Infect Immun 60, 1178–1186 (1992).

22. Hughes, C. V., Roseberry, C. A. & Kolenbrander, P. E. Isolation and characterization of coaggregation-defective mutants of *Veillonella atypica*. Archives of Oral Biology 35, S123– S125 (1990).

23. Zhou, P., Liu, J., Merritt, J. & Qi, F. A YadA-like autotransporter, Hag 1, in Veillonella atypica is a Multivalent Hemagglutinin Involved in Adherence to Oral Streptococci, Porphyromonas gingivalis, and Human Oral Buccal Cells. Mol Oral Microbiol 30, 269–279 (2015).

24. Zhou, P., Liu, J., Li, X., Takahashi, Y. & Qi, F. The Sialic Acid Binding Protein, Hsa, in Streptococcus gordonii DL1 also Mediates Intergeneric Coaggregation with Veillonella Species. PLOS ONE 10, e0143898 (2015).

25. Béchon, N. et al. Autotransporters Drive Biofilm Formation and Autoaggregation in the Diderm Firmicute Veillonella parvula. Journal of Bacteriology 202, (2020).

26. Nairn, B. L. et al. Uncovering Roles of Streptococcus gordonii SrtA-Processed Proteins in the Biofilm Lifestyle. Journal of Bacteriology 203, (2020).

27. Petersen, H. J. et al. Human Platelets Recognize a Novel Surface Protein, PadA, on Streptococcus gordonii through a Unique Interaction Involving Fibrinogen Receptor GPIIbIIIa. Infect Immun 78, 413–422 (2010).

28. Biswas, I., Jha, J. K. & Fromm, N. Shuttle expression plasmids for genetic studies in Streptococcus mutans. Microbiology (Reading) 154, 2275–2282 (2008).

29. Mutha, N. V. R. et al. Transcriptional profiling of coaggregation interactions between Streptococcus gordonii and Veillonella parvula by Dual RNA-Seq. Scientific Reports 9, 7664 (2019).

30. Jakubovics, N. S., Gill, S. R., Iobst, S. E., Vickerman, M. M. & Kolenbrander, P. E. Regulation of Gene Expression in a Mixed-Genus Community: Stabilized Arginine Biosynthesis in Streptococcus gordonii by Coaggregation with Actinomyces naeslundii. Journal of Bacteriology 190, 3646–3657 (2008).

31. Development of a Universal Fluorescent Probe for Gram-Positive Bacteria - Kwon - 2019 - Angewandte Chemie International Edition - Wiley Online Library. https://onlinelibrary.wiley.com/doi/full/10.1002/anie.201902537.

32. Takahashi, Y., Konishi, K., Cisar, J. O. & Yoshikawa, M. Identification and Characterization of hsa, the Gene Encoding the Sialic Acid-Binding Adhesin of Streptococcus gordonii DL1. Infection and Immunity 70, 1209–1218 (2002).

33. Haworth, J. A. et al. Concerted functions of Streptococcus gordonii surface proteins PadA and Hsa mediate activation of human platelets and interactions with extracellular matrix. Cellular Microbiology 19, e12667 (2017).

34. Béchon, N. et al. Capsular Polysaccharide Cross-Regulation Modulates Bacteroides thetaiotaomicron Biofilm Formation. mBio 11, e00729–20 (2020).

35. Bateman, A., Holden, M. T. G. & Yeats, C. The G5 domain: a potential N-acetylglucosamine recognition domain involved in biofilm formation. Bioinformatics 21, 1301–1303 (2005).

36. Corrigan, R. M., Rigby, D., Handley, P. & Foster, T. J. The role of Staphylococcus aureus surface protein SasG in adherence and biofilm formation. Microbiology (Reading) 153, 2435–2446 (2007).

37. Mutha, N. V. R. et al. Transcriptional responses of Streptococcus gordonii and Fusobacterium nucleatum to coaggregation. Mol Oral Microbiol 33, 450–464 (2018).

38. Kiliç, A. O. et al. Involvement of Streptococcus gordonii Beta-Glucoside Metabolism Systems in Adhesion, Biofilm Formation, and In Vivo Gene Expression. Journal of Bacteriology 186, 4246–4253 (2004).

39. Jakubovics, N. S. et al. Critical roles of arginine in growth and biofilm development by Streptococcus gordonii. Molecular Microbiology 97, 281–300 (2015).

40. Liu, J., Wu, C., Huang, I.-H., Merritt, J. & Qi, F. Differential response of Streptococcus mutans towards friend and foe in mixed-species cultures. Microbiology (Reading) 157, 2433– 2444 (2011).

41. Luppens, S. B. I. et al. Effect of Veillonella parvula on the antimicrobial resistance and gene expression of Streptococcus mutans grown in a dual-species biofilm. Oral Microbiology and Immunology 23, 183–189 (2008).

42. Chekli, Y. et al. Escherichia coli Aggregates Mediated by Native or Synthetic Adhesins Exhibit Both Core and Adhesin-Specific Transcriptional Responses. Microbiol Spectr 11, e0069023 (2023).

43. Sønderholm, M., et al. Pseudomonas aeruginosa Aggregate Formation in an Alginate Bead Model System Exhibits In Vivo-Like Characteristics. Applied and Environmental Microbiology 83, e00113–17 (2017).

44. Nadell, C. D., Drescher, K., Wingreen, N. S. & Bassler, B. L. Extracellular matrix structure governs invasion resistance in bacterial biofilms. ISME J 9, 1700–1709 (2015).

45. Burmølle, M. et al. Enhanced Biofilm Formation and Increased Resistance to Antimicrobial Agents and Bacterial Invasion Are Caused by Synergistic Interactions in Multispecies Biofilms. Appl Environ Microbiol 72, 3916–3923 (2006).

46. Lee, K. W. K. et al. Biofilm development and enhanced stress resistance of a model, mixed-species community biofilm. ISME J 8, 894–907 (2014).

47. Knapp, S. et al. Natural Competence Is Common among Clinical Isolates of Veillonella parvula and Is Useful for Genetic Manipulation of This Key Member of the Oral Microbiome. Front. Cell. Infect. Microbiol. 7, (2017).

48. Danne, C., Guérillot, R., Glaser, P., Trieu-Cuot, P. & Dramsi, S. Construction of isogenic mutants in Streptococcus gallolyticus based on the development of new mobilizable vectors. Research in Microbiology 164, 973–978 (2013).

49. Zhang, S., Zou, Z., Kreth, J. & Merritt, J. Recombineering in Streptococcus mutans Using Direct Repeat-Mediated Cloning-Independent Markerless Mutagenesis (DR-CIMM). Front Cell Infect Microbiol 7, 202 (2017).

50. Shields, R. C., Kaspar, J. R., Lee, K., Underhill, S. A. M. & Burne, R. A. Fluorescence Tools Adapted for Real-Time Monitoring of the Behaviors of Streptococcus Species. Appl. Environ. Microbiol. 85, (2019).

51. Cokelaer, T., Desvillechabrol, D., Legendre, R. & Cardon, M. ‘Sequana’: a Set of Snake-make NGS pipelines. Journal of Open Source Software 2, 352 (2017).

52. Köster, J. & Rahmann, S. Snakemake—a scalable bioinformatics workflow engine. Bioinformatics 28, 2520–2522 (2012).

53. Chen, S., Zhou, Y., Chen, Y. & Gu, J. fastp: an ultra-fast all-in-one FASTQ preprocessor. Bioinformatics 34, i884–i890 (2018).

54. Langmead, B. & Salzberg, S. L. Fast gapped-read alignment with Bowtie 2. Nat Methods 9, 357–359 (2012).

55. Liao, Y., Smyth, G. K. & Shi, W. featureCounts: an efficient general purpose program for assigning sequence reads to genomic features. Bioinformatics 30, 923–930 (2014).

56. Love, M. I., Huber, W. & Anders, S. Moderated estimation of fold change and dispersion for RNA-seq data with DESeq2. Genome Biology 15, 550 (2014).

57. Hartmann, R. et al. Quantitative image analysis of microbial communities with BiofilmQ. Nat Microbiol 6, 151–156 (2021).

