## Supplementary information for "Identification of *V. parvula* and *S. gordonii* adhesins mediating co-aggregation and its impact on physiology and mixed biofilm structure"

#### Table S1 : list of strains used in this study

| Strain | Description | Reference |
|---|---|---|
| ***Veillonella parvula* SKV38 strains** | | |
| SKV38 | WT | (Knapp et al. 2017) |
| *ΔvmaA* mutant | SKV38 *ΔFNLLGLLA_00032*::*tetM* | (Béchon et al. 2020) |
| *ΔvmaB* mutant | SKV38 *ΔFNLLGLLA_00335*::*tetM* | (Béchon et al. 2020) |
| *ΔvmaC* mutant | SKV38 *ΔFNLLGLLA_00581*::*tetM* | (Béchon et al. 2020) |
| Δ*vtaA* mutant | SKV38 *ΔFNLLGLLA_00516*::*tetM* | (Béchon et al. 2020) |
| *ΔvtaB* mutant | SKV38 *ΔFNLLGLLA_00034*::*tetM* | (Béchon et al. 2020) |
| *ΔvtaC* mutant | SKV38 *ΔFNLLGLLA_00038*::*tetM* | This study |
| *ΔvtaD* mutant | SKV38 *ΔFNLLGLLA_00044*::*tetM* | This study |
| *ΔvtaE* mutant | SKV38 *ΔFNLLGLLA_00045*::*tetM* | This study |
| *ΔvtaF* mutant | SKV38 *ΔFNLLGLLA_00046*::*tetM* | This study |
| *ΔvtaG* mutant | SKV38 *ΔFNLLGLLA_00098*::*tetM* | (Béchon et al. 2020) |
| *ΔvtaH* mutant | SKV38 *ΔFNLLGLLA_00099*::*tetM* | (Béchon et al. 2020) |
| *ΔvtaI* mutant | SKV38 *ΔFNLLGLLA_01790*::*tetM* | (Béchon et al. 2020) |
| *ΔvtaA ΔvtaC-F* mutant | SKV38 *ΔFNLLGLLA_ 00516*::*catP ΔFNLLGLLA_00036-46*::*tetM* | (Béchon et al. 2020) |
| *ΔvtaC-F* mutant | SKV38 *ΔFNLLGLLA_00036-46*::*tetM* | (Béchon et al. 2020) |
| $P_{Tet}$ -*vtaA* mutant | SKV38 *catP-Term(fdx)*-P*tet-FNLLGLLA_00516* | (Béchon et al. 2020) |
| $P_{Tet}$ -*vtaE* mutant | SKV38 *ermE-Term(fdx)*-P*tet-FNLLGLLA_00045* | This study |
| Δ*vtaE* Δ*vtaC* mutant | SKV38 *ΔFNLLGLLA_ 00045*::kanR *ΔFNLLGLLA_00043*::*tetM* | This study |
| Δ*vtaD-E* mutant | SKV38 *ΔFNLLGLLA_00044-45*::*tetM* | This study |
| Δ*vtaE* Δ*vtaF* mutant | SKV38 *ΔFNLLGLLA_ 00045*::kanR *ΔFNLLGLLA_00046*::*tetM* | This study |
| Δ*vtaD-E* Δ*vtaA* | SKV38 *ΔFNLLGLLA_00044-45*::*tetM* ΔFNLLGLLA_ 00516::catP | This study |
| ***Streptococcus gordonii* DL1** | | |
| DL1 | WT | (Nairn et al. 2020) |
| ΔsrtA | DL1 ΔSGO_1230 | (Nairn et al. 2020) |
| Δ1487 | *DL1 ΔSGO_1487* | (Nairn et al. 2020) |
| ΔabpB | DL1 Δ*SGO*_0162 | (Nairn et al. 2020) |
| ΔcshB | DL1 ΔSGO_1148 | (Nairn et al. 2020) |
| Δhsa | DL1 ΔSGO_0966 | (Nairn et al. 2020) |
| ΔsspAB | DL1 Δ*SGO_0210-0211* | (Nairn et al. 2020) |

| | | |
|---|---|---|
| Δ0107 | DL1 ΔSGO_0107 | (Nairn et al. 2020) |
| Δ0208 /Δ*endoD* | DL1 ΔSGO_0208 | (Nairn et al. 2020) |
| Δ0316 | DL1 Δ*SGO_0316* | (Nairn et al. 2020) |
| Δ0317 | DL1 ΔSGO_0317 | (Nairn et al. 2020) |
| Δ0385 | DL1 ΔSGO_0385 | (Nairn et al. 2020) |
| Δ0388 | DL1 ΔSGO_0388 | (Nairn et al. 2020) |
| Δ0405/Δ*strH* | DL1 ΔSGO_0405 | (Nairn et al. 2020) |
| Δ0430 | DL1 ΔSGO_0430 | (Nairn et al. 2020) |
| Δ0707 | DL1 ΔSGO_0707 | (Nairn et al. 2020) |
| Δ0854/ΔcshA | DL1 Δ*SGO_0854* | (Nairn et al. 2020) |
| Δ0890 | DL1 ΔSGO_0890 | (Nairn et al. 2020) |
| Δ1182 / ΔpavB | DL1 ΔSGO_1182 | (Nairn et al. 2020) |
| Δ1247 / ΔNt5e | DL1 Δ*SGO_1247* | (Nairn et al. 2020) |
| Δ1415 | DL1 Δ*SGO_1415* | (Nairn et al. 2020) |
| Δ1435 | DL1 ΔSGO_1435 | (Nairn et al. 2020) |
| Δ1486 / ΔBgaA | DL1 Δ*SGO_1486* | (Nairn et al. 2020) |
| ΔabpA | DL1 ΔSGO_2105 | (Nairn et al. 2020) |
| Δ1650 | DL1 ΔSGO_1650 | (Nairn et al. 2020) |
| Δ1651 | DL1 ΔSGO_1651 | (Nairn et al. 2020) |
| Δ1727 | DL1 ΔSGO_1727 | (Nairn et al. 2020) |
| Δ2004 | DL1 ΔSGO_2004 | (Nairn et al. 2020) |
| Δ*padA* | DL1 Δ*SGO_2005* | (Nairn et al. 2020) |
| IDFC2-visA | DL1 IDFC2-*SGO_2004* | This study |
| $P_{Tet}$-visA | DL1 pTeT-pVeg(RBS)-*SGO_2004* | This study |
| Δ*padA* $P_{Tet}$ -visA | DL1 Δ*padA* pTeT-pVeg(RBS)-*SGO_2004* | This study |
| ***Escherichia coli* strains** | | |
| DH5α | WT | Laboratory collection |
| Bl21(DE3) pDIA17 | WT | (Munier et al. 1991) |
| Bl21(DE3) pDIA17 pET22b-VisA-HIS | pET22b-VisA-His | This study |
| **Other bacteria** | | |
| *S. oralis CIP103216* | WT | CIP Institut Pasteur |

| | | |
|---|---|---|
| *S. parasanguinis CIP104372T* | WT | CIP Institut Pasteur |
| *S. gordonii CIP105038* | WT | CIP Institut Pasteur |
| *S. gordonii CIP105258T* | WT | CIP Institut Pasteur |
| *S. gordonii NCTC7868* | WT | CIP Institut Pasteur |
| *S. mitis CIP104996* | WT | CIP Institut Pasteur |
| *S. mitis CIP104997* | WT | CIP Institut Pasteur |
| *A. oris CIP102340* | WT | CIP Institut Pasteur |
| *F. nucleatum CIP101130T* | WT | CIP Institut Pasteur |
| *S. mutans NG8* | WT | Justin Meritt(Lee et al. 1989) |
| *S. mutans UA140* | WT | (Switalski and Butcher 1994) |
| *S. mutans UA159* | WT | (Ajdić et al. 2002)8 |
| *S. mutans CBSm8* | WT | Justin Meritt |
| *S. mutans CBSm38* | WT | Justin Meritt |

#### Table S2 : primers used in this study

| name | Sequence (5'->3') | construction |
|---|---|---|
| 0045-5F | actaaagatggcctcaac | Upstream of Ptet-VtaE/ΔvtaE-KanR |
| 5R_pTet_VtaE | tcaacacactcttaagtttgcttcctcatcttactccttgcataaaac | Upstream of Ptet-VtaE |
| JW264 | gaagcaaacttaagagtgtgttga | Ptet-ery cassette |
| K7ind-ptet-cat-R | aaaattttctcctttactgcagg | Ptet-ery cassette |
| 3F_ptet_VtaE | cctgcagtaaaggagaaaattttatgaaccgaatttttaaagtaatctg | Downstream of Ptet-VtaE |
| 0044_3R | ccgcctttagatgccttc | Downstream of Ptet-VtaE |
| 5R_delta_vtaE_kana | cactcgcctgcattttactcatcttactccttgcataaaac | Upstream of ΔvtaE-KanR |
| F_kana | TAAAATGCAGGCGAGTGAAGaaaattttc | KanR cassette |
| R_kana | TAAAACAATTCATCCAGTAAAATATAATATTTTATTTTCTCCC | KanR cassette |
| 3F_delta_vtaE_kana | ATTTTACTGGATGAATTGTTTTAGttcacatctgtttcatcttatattc | Downstream of ΔvtaE-KanR |
| 0045_3R | gccaaggagttaaaaccc | Downstream of ΔvtaE-KanR |
| 5F_SGO2004prom | gattccgaataaggctactg | Upstream of pTet-SGO2004/IDFC2-SGO2004 |
| 5R_SGO2004prom_IDFC | GATGAGCGTTATCGTGGCTCttaatgctttggttttcttctc | Upstream of IDFC2-SGO2004 |
| F_IDFC_sgordonii | GAGCCACGATAACGCTCATC | IDFC cassette |

| | | |
|---|---|---|
| R_IDFC_sgordonii | ttatttcctcccgttaaataatagataac | IDFC cassette |
| 3F_2004prom_IDFC | GTTATCTATTATTTAACGGGAGGAAATAAatggatagaaaaaagttaattaagttag | Downstream of IDFC2-SGO2004 |
| 3R_2004_prom | cttggtaacgagcttcatac | Downstream of pTet-SGO2004/IDFC2-SGO2004 |
| 5R_SGO2004_pTet | gcagaattcgcccttaagtcttaatgctttggttttcttctc | Upstream of pTet-SGO2004 |
| F_pTet_promoter | gacttaagggcgaattctgc | Ptet-pVEG_RBS cassette |
| R_pTet_pVEG_RBS | GTTTGTCCTCCTTATTAGTTAATCAagctcagatctgttaacgcta | Ptet-pVEG_RBS cassette |
| 3F_2004_pTET_pvegRBS | TGATTAACTAATAAGGAGGACAAACatggatagaaaaaagttaattaagttag | Downstream of pTet-SGO2004 |
| 5F_delta_padA | gagcggatcagtatatcagtg | Upstream of ΔpadA pTet-SGO2004 |
| 5R_delta_padA_pTetSGO | gcagaattcgcccttaagtcgctattttaaagcctatataaatagtag | Upstream of ΔpadA pTet-SGO2004 |
| F_SGO2004_pet22b | CTTTAAGAAGGAGATATACATatgcaaccagagattacgactcag | SGO_2004(aa136-698) for pET22b |
| R_SGO2004_pet22b | TCAGTGGTGGTGGTGGTGGTGttccttagtaccaatttcaactac | SGO_2004(aa136-698) for pET22b |
| pET22b_LINEAR_F | CACCACCACCACCACCACT | linearization of pET22b |
| pET22Bb_LINEAR_R | ATGTATATCTCCTTCTTAAAGTTAAACAAAATTATTTC | linearization of pET22b |

#### Supplementary figures

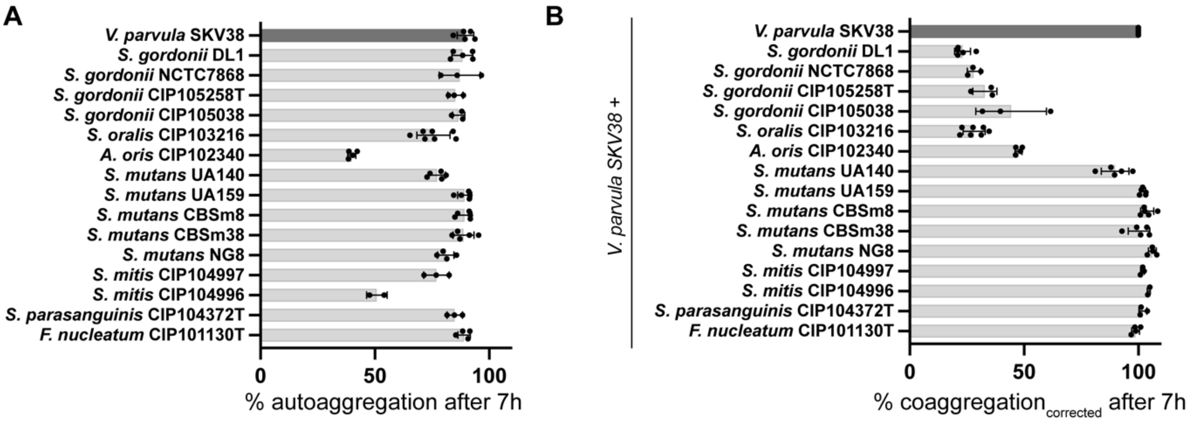


**Figure S1: Auto-aggregation and co-aggregation between *V. parvula* SKV38 and various members of the dental plaque.**

(A) Auto-aggregation of *V. parvula* SKV38 and various members of the dental plaque after 7h, as measured by the % of decrease of optical density between 0 and 7h. SD and single points for 3-5 replicates are shown. (B) Co-aggregation from independent cultures corrected for auto-aggregation of *V. parvula* SKV38 with various members of the dental plaque after 7h, as measured by the % of decrease of optical density between 0 and 7h. SD and single points for 3-5 replicates are shown. Correction was applied using the following formula:

$$Coaggregation_{corrected} = Coaggregation + 100 - \left(\frac{autoaggregation_{Vp} + autoaggregation_{partner}}{2}\right)$$

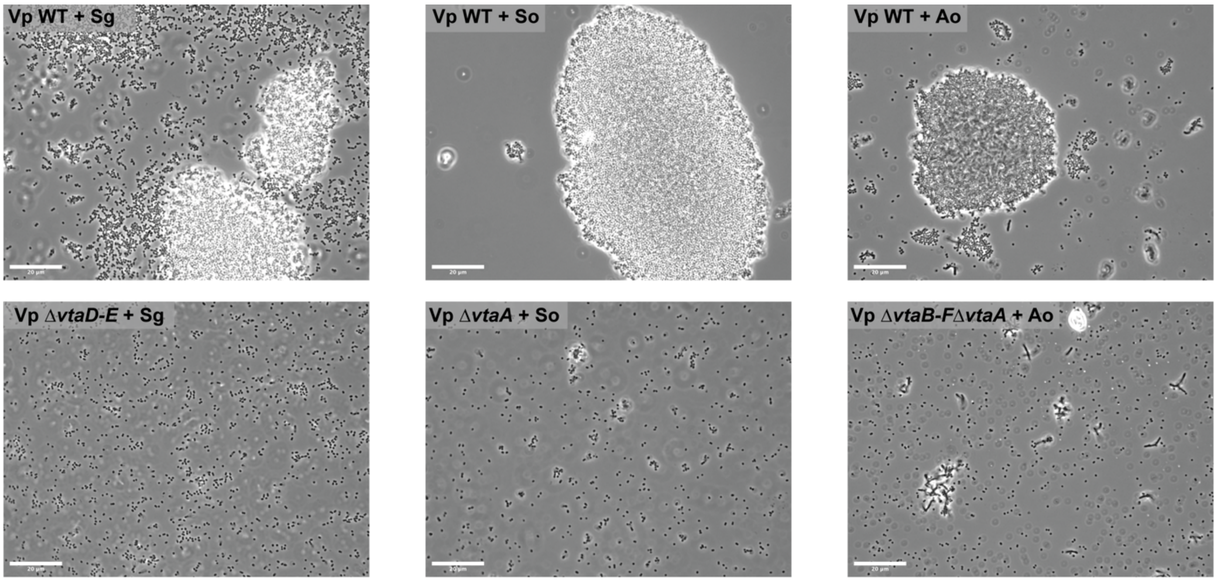


**Figure S2: Light microscopy images of co-aggregates between *V. parvula* SKV38 and *S. oralis*, *S. gordonii* and *A. oris*.**

Brightfield images of aggregates after 7h of co-aggregation. 3 µl of bacteria solution were sampled at the bottom of a cuvette used for an aggregation assay, mounted on a slide and observed under the microscope. Scale bar is 20 µm. Sg: *S. gordonii*, So: *S. oralis*, Ao: *A. oris*.

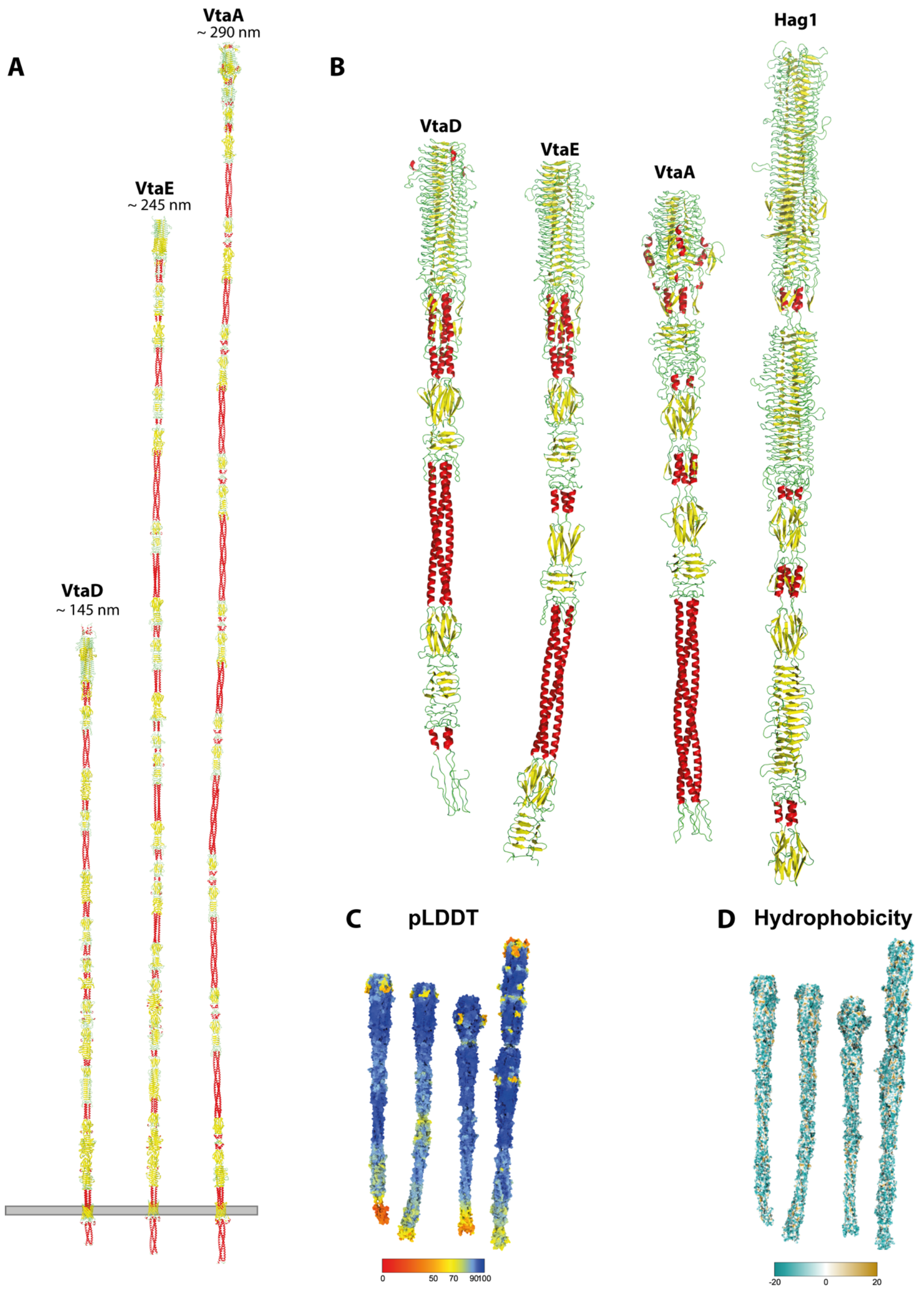


**Figure S3: Comparison of the predicted structures of *V. parvula* VtaD, VtaE, VtaA and *V. atypica* Hag1.**

(A) Model predictions of the full length trimeric VtaD, VtaE and VtaA. Models were obtained by predicting 400-700 residues overlapping fragments of each adhesin as trimers using AlphaFold 2.3(Evans et al. 2022) locally. Fragments were then assembled in PyMOL using the command Super. (B-C-D) Comparison of the head structure, pLDDT (per-residue model confidence score, higher values mean higher confidence in the prediction) and hydrophobicity of VtaD, VtaE, VtaA and Hag1. Images, pLDDT coloring and hydrophobicity calculation were done in ChimeraX(Pettersen et al. 2021).

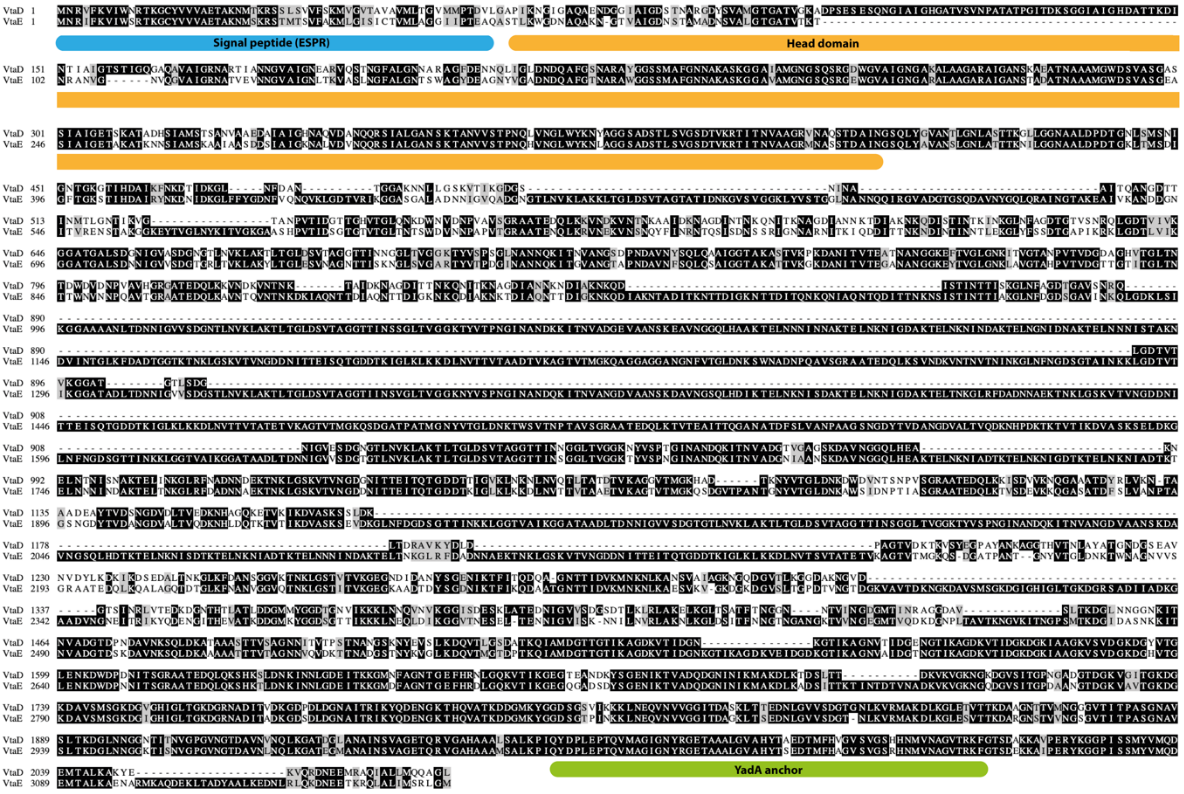


**Figure S4: Sequence alignment between *V. parvula* VtaE and VtaD.**
Both protein sequences were aligned using the Emboss Water algorithm(Madeira et al. 2022). The ESPR sequence is highlighted in blue, the head section of the adhesins in orange and the YadA anchor in green. Identical amino acid sequences are highlighted in black, similar in gray and divergent in white.

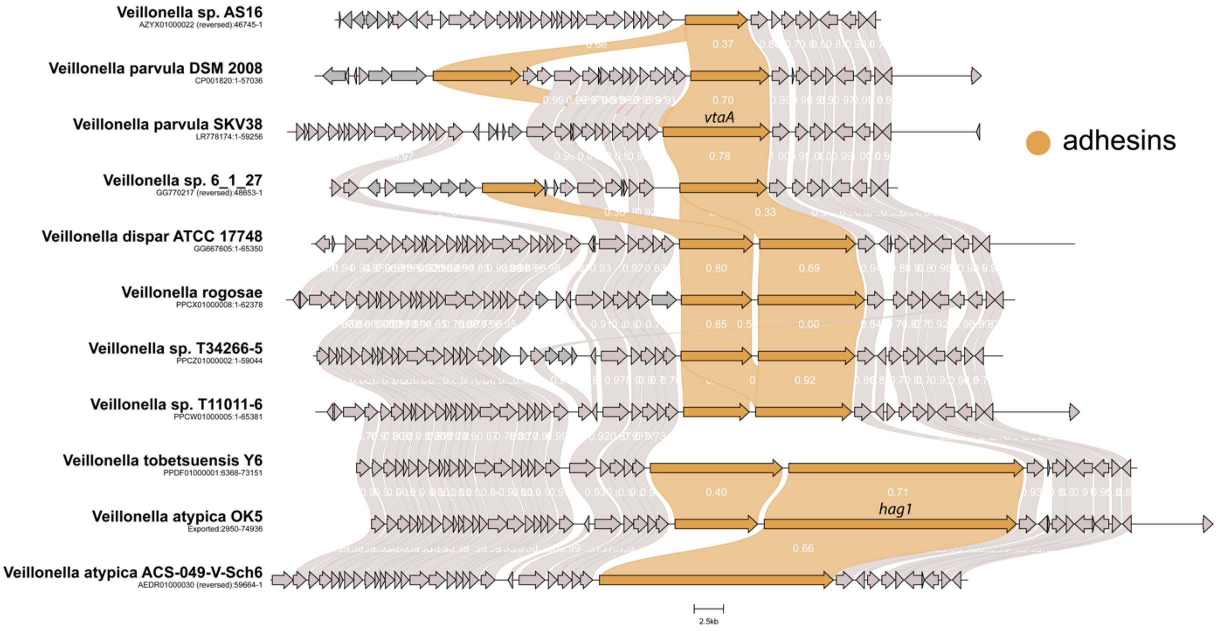


**Figure S5: Variations at the *vtaA* locus in *Veillonella*.**
A local subset of Veillonella genomes was searched for presence of the *vtaA* neighboring MATE efflux transporting using the HMM profile PF01554.21. Neighboring regions were extracted and aligned using clinker(Gilchrist and Chooi 2021). Trimeric autotransporters are colored in brown.

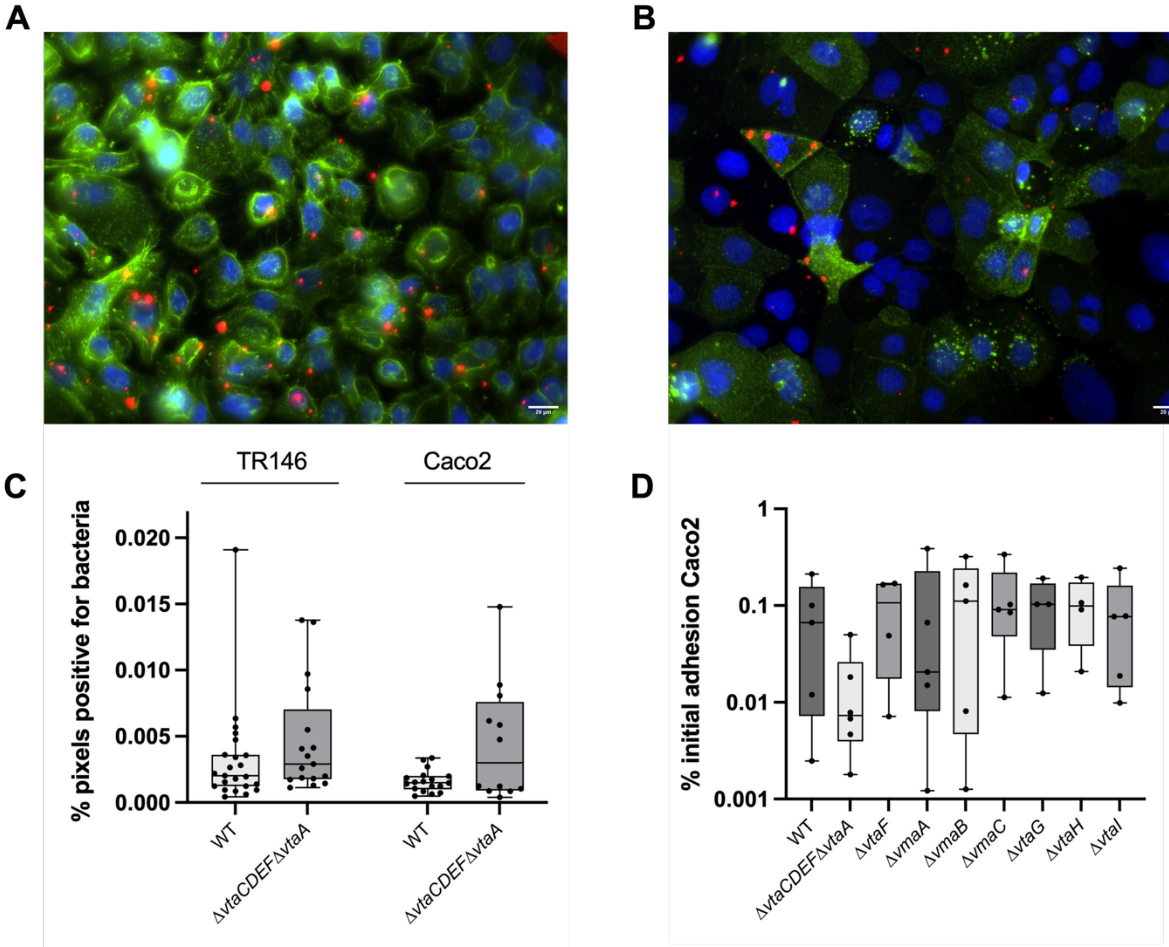


**Figure S6: *V. parvula* presents a limited adhesion to host cells**.
Representative image of *V. parvula* SKV38 WT adhesion on buccal TR146 (A) and intestinal Caco-2 epithelial cells (B) after 30min. Bacteria are labelled in red, human cell surface labelled in green and nuclei in blue. Scale bar is 20 µm. (C) Quantification of the adhesion shown in A-B, (N=2) as the percentage of pixels positive for bacteria in the Z-stack (See supplementary methods). No statistical differences were observed between the different groups using a Kruskal-Wallis test. (D) Ratio of adhering bacteria compared to the total amount of bacteria after 2h of incubation on Caco-2 cells for *V. parvula* WT and different adhesin mutants (N=4 to 5).

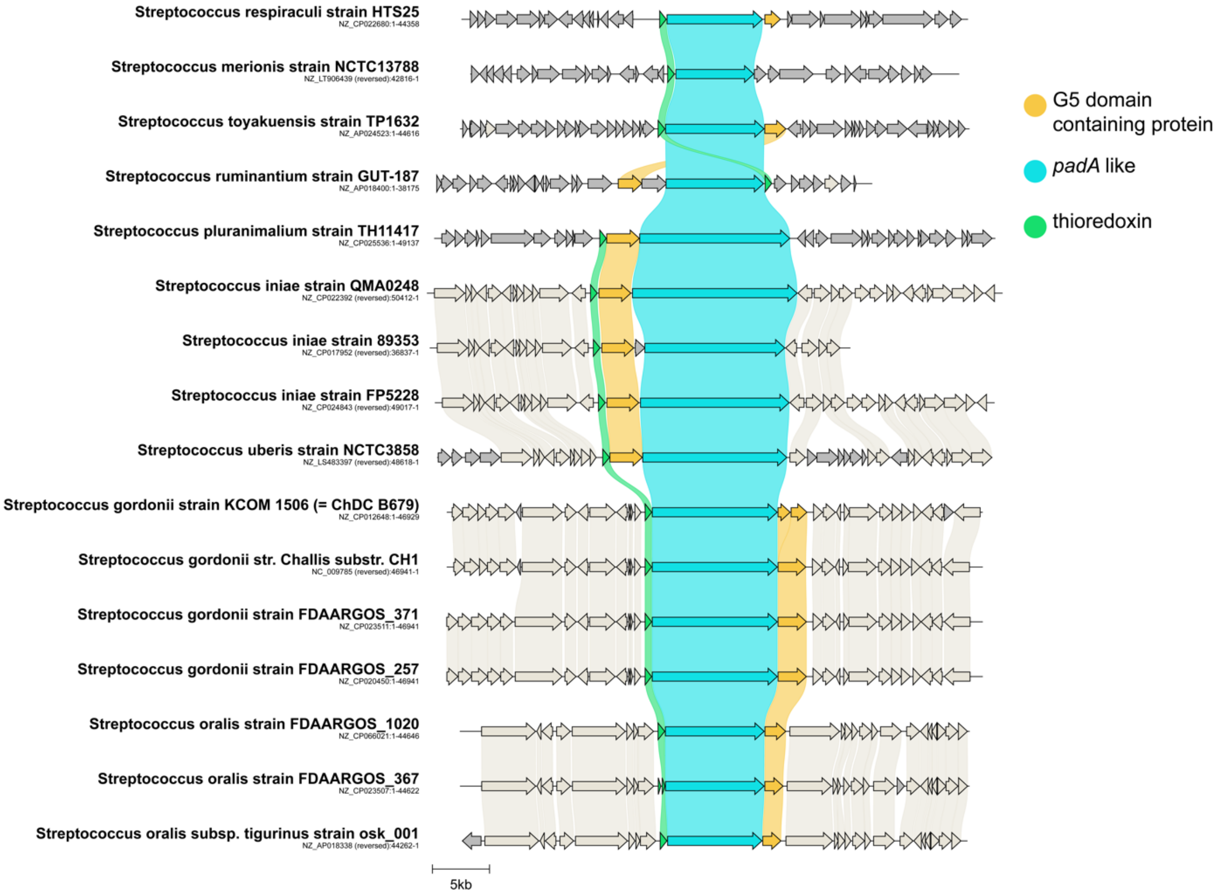


#### Figure S7: Conservation of the *padA* operon in distant streptococci.

A local subset of Streptococcus genomes was searched for presence of PadA using a custom HMM profile. Neighboring regions were extracted and aligned using clinker(Gilchrist and Chooi 2021). *padA* homologs are colored in blue, neighboring G5 domain containing proteins in yellow and thioredoxins in green.

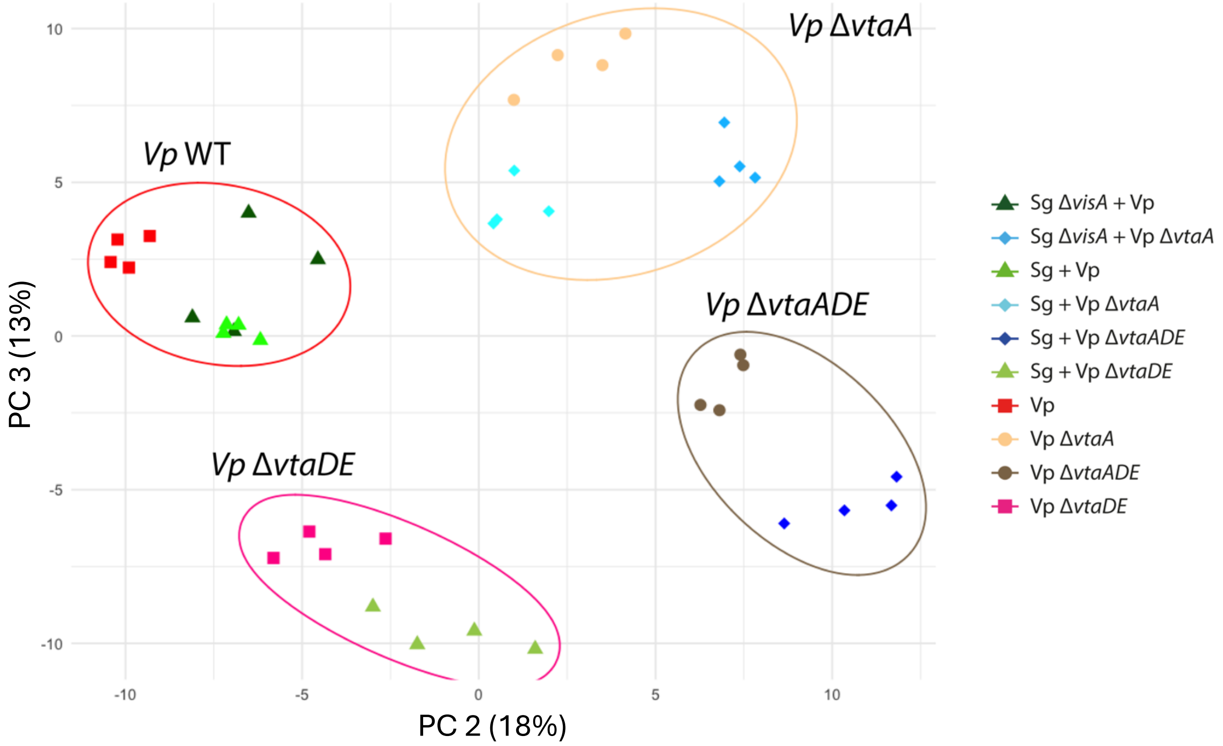


**Figure S8: 2nd and 3rd dimension of principal component analysis (PCA) of all *V. parvula* samples.**

These experiments correspond to 4 biological replicates for 10 conditions. Colors and shape represent the different conditions. The four circles separate samples based on *Veillonella* strains.

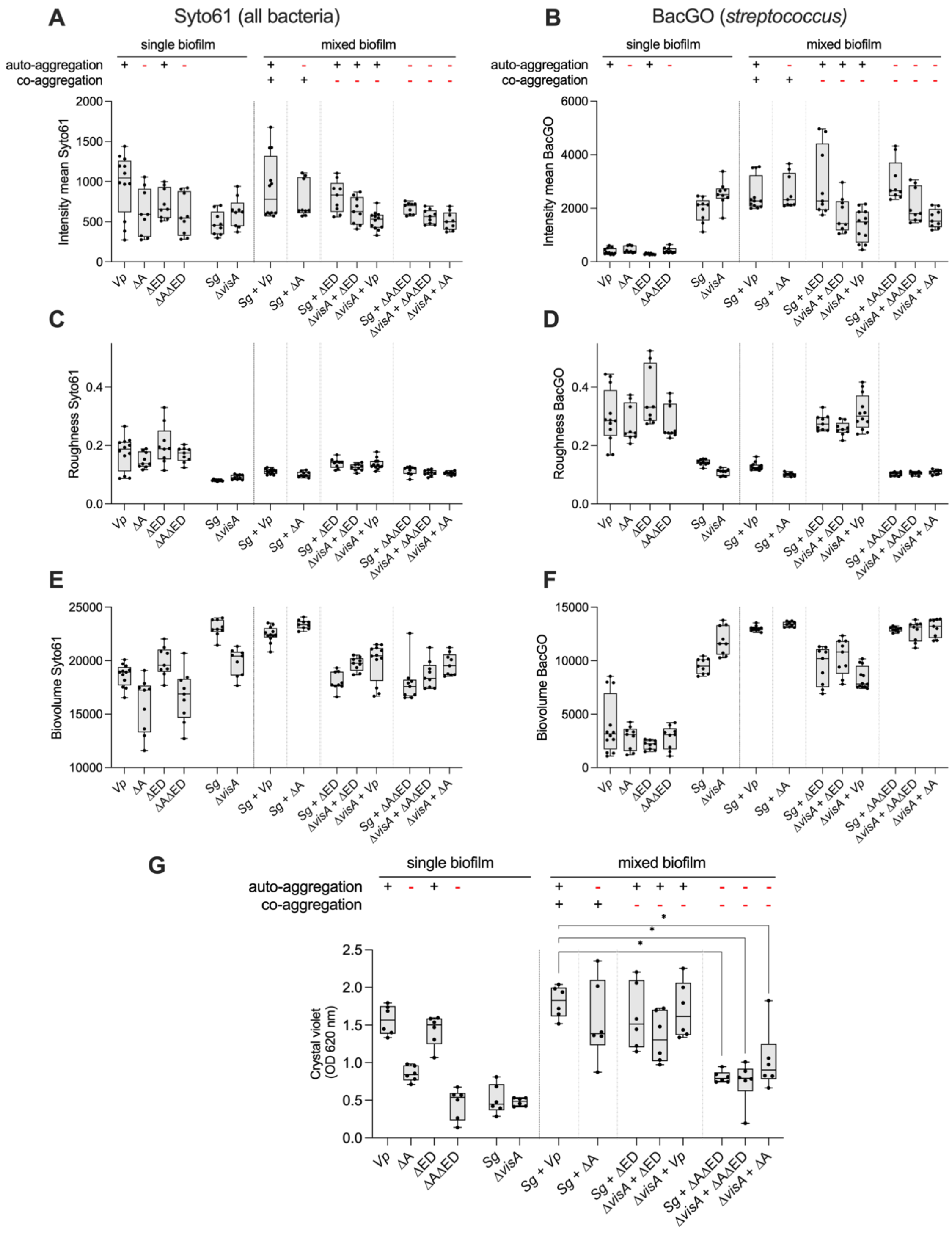

**Figure S9**: **Biofilms calculated parameters**.

Average intensity of each channel (A-B), biofilm roughness (C-D) and the Biofilm volume (E-F) were calculated for the BacGO dye (*Streptococcus*) and Syto61 dye (all bacteria) using BiofilmQ 1.1. Each point (9-12 per condition) represents the average roughness measurement of four images per well. Experiment done in three biological independent replicates. G) Estimations of biofilm formation in BHIP medium in a 96-well plate by crystal violet staining. Min-max box plots for 6 biological replicates. Each replicate is the mean of two technical replicates. A Kurskal-wallis test comparing all mixed biofilms conditions to *Sg -V. parvula* followed by a Benjamini, Krieger and Yekutieli correction for multiple testing was used. Significant P values (<0.05) are indicated by an asterisk. For all plots, *Vp* is *V. parvula*, ΔA is *V. parvula* Δ*vtaA*, ΔED is *V. parvula* Δ*vtaD-vtaE*, *Sg* is *S. gordonii* and Δ*visA* is *S. gordonii* Δ*visA*. Presence (or absence) of auto- and co- aggregation is indicated by the + (or -) symbols.

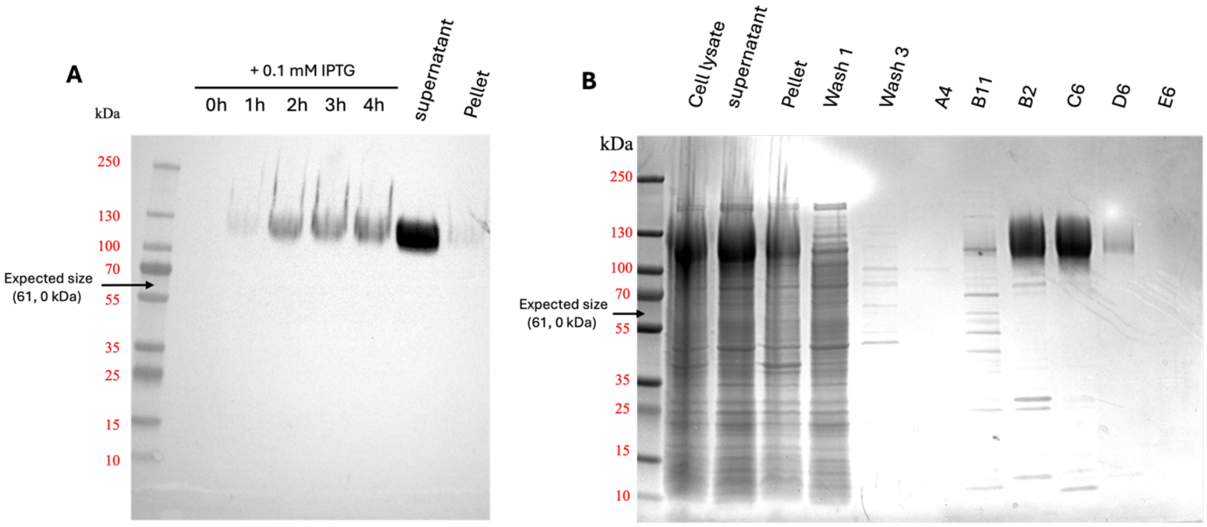


**Figure S10: Purification of VisA$_{G5}$**

A) Anti His-Tag Western blot on *E. coli* whole cell lysates at different times after protein production induction by IPTG. Supernatant and pellet correspond to the different fractions of a 4h induced *E. coli* lysate by BugBuster Protein Extraction Reagent. B) Whole protein gel stained with SimplyBlue SafeStain corresponding to the different steps of protein purifications. 60 fractions (A1 to E12) were eluted and collected among which selected fractions were deposited on the presented gel. Fractions B6 to D10 were pooled together to yield the final purified protein. The arrows indicate the expected size of a monomer of the purified protein.

### Supplementary methods

**Biofilm formation in 96-well microtiter plates.**

Overnight cultures of *S. gordonii* strains in BHI and *V. parvula strains* in BHIL medium were diluted to an $OD_{600}$ of 0.05 and transferred to two flat-bottom TPP 96-well plates (Dustscher), adding 150 µL per well. After 24 h of static incubation in anaerobia, supernatant was removed, and one of the two plates was resuspended by pipetting to measure $OD_{600}$ using a Tecan Infinite-M200-Pro spectrophotometer. The other plate was used for coloration. Biofilms were fixed with 150 µL Bouin solution (HT10132; Sigma-Aldrich) for 15 min. Bouin solution was removed by inversion, and the biofilms were washed once in water. The biofilms were stained with 150 µL of 1% crystal violet (V5265; Sigma-Aldrich) for 15 min without shaking and then washed in water three times and left to dry. All washes were made by flicking the plate. After drying the plate, crystal violet was dissolved with 200 µL absolute ethanol and $OD_{620}$ was measured.

#### Culture of the human epithelial cells

Caco-2 and TR146 (CancerTools Ref 151425) cells were maintained in DMEM Glutamax (Gibco) supplemented with 10% of heat-inactivated serum and non-essential amino acids (Gibco). Caco2 cells were grown at 37°C in a 10% CO2 incubator and TR146 cells in a 5% CO2 incubator. Contamination to mycoplasma was regularly tested using PCR test and HEK hTLR2 (Invivogen).

#### Adhesion of *V. parvula* adhesin mutants on Caco-2 cells

The day prior each experiment, fresh culture of *V. parvula* strains were inoculated and Caco-2 cells were seeded at 0.4 x $10^6$ cells per well in 12 wells plates (TPP). On the day of experiment, Caco-2 medium was removed and changed to DMEM without serum and placed in the anaerobic chamber (Jacomex) at least 1h before the adhesion. In the anaerobic chamber, *V. parvula* strains were centrifuged at 5000g for 10min at room temperature, washed with PBS, centrifuged again before resuspension in DMEM without serum at OD=1. For the adhesion, 10 µL of each strain/well were added and incubated for 2h at 37°C. After incubation, the supernatant was removed from each well, and Caco-2 cells were washed 3x with 1 mL PBS before adding 50µL of Trypsin-EDTA 0.25% (Gibco) and 15min incubation at 37°C. To resuspend Caco-2 cell lysate, 50 µL of PBS was added and homogenised by pipetting. Serial dilutions for each mutant were performed from $10^{-1}$ to $10^{-7}$ in PBS and plated in duplicate on SK agar plates and left for incubation at 37°C for 2 days.

#### Comparison of adhesion on buccal and intestinal epithelial cells by microscopy

Two days before the experiment, Caco-2 and TR146 cells were seeded at 0.4x $10^6$ cells per well on glass cover slips in 24 wells plate. One day prior experiment the bacterial culures were refreshed. On the day of experiment, 1 mL of culture of each strain were harvested by centrifugation at 5000g for 8min. Bacteria were resuspended in PBS at OD=1 and 0.5µl of Cellbrite 640 (Biotium) was added to 500 µl of bacterial and incubate 30min at 37°C with shaking at 300rpm. To wash the dye, 1 mL of PBS was added before centrifugation at 5000 g for 8 min and bacteria were resuspended in PBS. To dislodge aggregates, bacteria were sonicated 2 x 10s in a water bath sonicator and 100 µM of each bacteriam culture was added to each epithelial cell well and let to adhere for 30min at 37°C. After the incubation, the wells were washed 3 times with PBS before fixation with PFA 4% for 40min at room temperature. After fixation and, cells were washed with PBS and cell surface was stained with WGA-555 10 µg/mL (Invivogen) for 45min, washed with PBS and stained with DAPI 2 µg/mL for another 10 min, all these steps were done at room temperature. After 2 final washes with PBS, cover slips were mounted with Prolong Gold and cured overnight.

A minimum of 5 fields were taken with Z stack for each cover slip, with 40X objective on a IX81 Olympus microscope. Each image was chosen on the DAPI channel and acquired on the 3 channels (DAPI, Cy3, Cy5). Images were quantified with Fiji by duplicating each image on the Cy5 channel and applying an automated threshold (Otsu method). Pixels occupied by the bacteria for each image of the stack were summed as well as the total number of pixels and ratios were calculated for each field.

**Supplementary references**


Ajdić, Dragana, William M. McShan, Robert E. McLaughlin, Gorana Savić, Jin Chang, Matthew B. Carson, Charles Primeaux, et al. 2002. “Genome Sequence of Streptococcus Mutans UA159, a Cariogenic Dental Pathogen.” *Proceedings of the National Academy of Sciences* 99 (22): 14434–39. https://doi.org/10.1073/pnas.172501299.

Béchon, Nathalie, Alicia Jiménez-Fernández, Jerzy Witwinowski, Emilie Bierque, Najwa Taib, Thomas Cokelaer, Laurence Ma, Jean-Marc Ghigo, Simonetta Gribaldo, and Christophe Beloin. 2020. “Autotransporters Drive Biofilm Formation and Autoaggregation in the Diderm Firmicute Veillonella Parvula.” *Journal of Bacteriology* 202 (21). https://doi.org/10.1128/JB.00461-20.

Evans, Richard, Michael O’Neill, Alexander Pritzel, Natasha Antropova, Andrew Senior, Tim Green, Augustin Žídek, et al. 2022. “Protein Complex Prediction with AlphaFold-Multimer.” bioRxiv. https://doi.org/10.1101/2021.10.04.463034.

Gilchrist, Cameron L M, and Yit-Heng Chooi. 2021. “Clinker & Clustermap.Js: Automatic Generation of Gene Cluster Comparison Figures.” *Bioinformatics* 37 (16): 2473–75. https://doi.org/10.1093/bioinformatics/btab007.

Knapp, Steven, Clint Brodal, John Peterson, Fengxia Qi, Jens Kreth, and Justin Merritt. 2017. “Natural Competence Is Common among Clinical Isolates of Veillonella Parvula and Is Useful for Genetic Manipulation of This Key Member of the Oral Microbiome.” *Frontiers in Cellular and Infection Microbiology* 7. https://doi.org/10.3389/fcimb.2017.00139.

Lee, S F, A Progulske-Fox, G W Erdos, D A Piacentini, G Y Ayakawa, P J Crowley, and A S Bleiweis. 1989. “Construction and Characterization of Isogenic Mutants of Streptococcus Mutans Deficient in Major Surface Protein Antigen P1 (I/II).” *Infection and Immunity* 57 (11): 3306–13. https://doi.org/10.1128/iai.57.11.3306-3313.1989.

Madeira, Fábio, Matt Pearce, Adrian R N Tivey, Prasad Basutkar, Joon Lee, Ossama Edbali, Nandana Madhusoodanan, Anton Kolesnikov, and Rodrigo Lopez. 2022. “Search and Sequence Analysis Tools Services from EMBL-EBI in 2022.” *Nucleic Acids Research* 50 (W1): W276–79. https://doi.org/10.1093/nar/gkac240.

Munier, Hélène, Anne-Marie Gilles, Philippe Glaser, Evelyne Krin, Antoine Danchin, Robert Sarfati, and Octavian Bârzu. 1991. “Isolation and Characterization of Catalytic and Calmodulin-Binding Domains of Bordetella Pertussis Adenylate Cyclase.” *European Journal of Biochemistry* 196 (2): 469–74. https://doi.org/10.1111/j.1432-1033.1991.tb15838.x.

Nairn, Brittany L., Grace T. Lee, Ashwani K. Chumber, Patrick R. Steck, Mahmoud O. Mire, Bruno P. Lima, and Mark C. Herzberg. 2020. “Uncovering Roles of Streptococcus Gordonii SrtA-Processed Proteins in the Biofilm Lifestyle.” *Journal of Bacteriology* 203 (2). https://doi.org/10.1128/JB.00544-20.

Pettersen, Eric F., Thomas D. Goddard, Conrad C. Huang, Elaine C. Meng, Gregory S. Couch, Tristan I. Croll, John H. Morris, and Thomas E. Ferrin. 2021. “UCSF ChimeraX: Structure Visualization for Researchers, Educators, and Developers.” *Protein Science: A Publication of the Protein Society* 30 (1): 70–82. https://doi.org/10.1002/pro.3943.

Switalski, L. M., and W. G. Butcher. 1994. “An in Vitro Model for Adhesion of Bacteria to Human Tooth Root Surfaces.” *Archives of Oral Biology* 39 (2): 155–61. https://doi.org/10.1016/0003-9969(94)90111-2.
